# Detection and classification of hard and soft sweeps from unphased genotypes by multilocus genotype identity

**DOI:** 10.1101/281063

**Authors:** Alexandre M. Harris, Nandita R. Garud, Michael DeGiorgio

## Abstract

Positive natural selection can lead to a decrease in genomic diversity at the selected site and at linked sites, producing a characteristic signature of elevated expected haplotype homozygosity. These selective sweeps can be hard or soft. In the case of a hard selective sweep, a single adaptive haplotype rises to high population frequency, whereas multiple adaptive haplotypes sweep through the population simultaneously in a soft sweep, producing distinct patterns of genetic variation in the vicinity of the selected site. Measures of expected haplotype homozygosity have previously been used to detect sweeps in multiple study systems. However, these methods are formulated for phased haplotype data, typically unavailable for nonmodel organisms, and may have reduced power to detect soft sweeps due to their increased genetic diversity relative to hard sweeps. To address these limitations, we applied the H12 and H2/H1 statistics of Garud et al. [2015] to unphased multilocus genotypes, denoting them as G12 and G2/G1. G12 (and the more direct expected homozygosity analogue to H12, denoted G123) has comparable power to H12 for detecting both hard and soft sweeps. G2/G1 can be used to classify hard and soft sweeps analogously to H2/H1, conditional on a genomic region having high G12 or G123 values. The reason for this power is that under random mating, the most frequent haplotypes will yield the most frequent multilocus genotypes. Simulations based on parameters compatible with our recent understanding of human demographic history suggest that expected homozygosity methods are best suited for detecting recent sweeps, and increase in power under recent population expansions. Finally, we find candidates for selective sweeps within the 1000 Genomes CEU, YRI, GIH, and CHB populations, which corroborate and complement existing studies.

## Introduction

Positive natural selection is the process by which an advantageous genetic variant rises to high frequency in a population, thereby reducing site diversity and creating a tract of elevated expected homozygosity and linkage disequilibrium (LD) surrounding that variant [Sabeti et al., 2002]. As beneficial alleles increase to high frequency in a population, the signature of a selective sweep emerges, which we can characterize from the number of haplotypes involved in the sweep [Maynard Smith and Haigh, 1974, Schweinsberg and Durrett, 2005, Hermisson and Pennings, 2017]. A hard sweep is an event in which a single haplotype harboring a selectively advantageous allele rises in frequency, while in a soft sweep, multiple haplotypes harboring advantageous mutations can rise in frequency simultaneously. Thus, selective sweeps represent a broad and non-homogenous spectrum of genomic signatures. A selective event that persists until the beneficial allele reaches fixation is a *complete* sweep, while a *partial* sweep is one in which the selected allele does not reach fixation. Consequently, expected haplotype homozygosity surrounding the selected site is greatest once the selected allele has fixed and before recombination and mutation break up local LD [Przeworski, 2002].

Two modes of soft sweeps have been proposed across three seminal papers, consisting of sweeps from standing genetic variation that becomes beneficial in a changing environment, or new recurrent *de novo* adaptive mutations [Hermisson and Pennings, 2005, Pennings and Hermisson, 2006a, b], and these can be complete and partial as well. Unlike hard sweeps, where haplotypic diversity is decreased, in a soft sweep, haplotypic diversity remains, since multiple haplotypes carrying the adaptive allele rise to high frequency. [Przeworski et al., 2005, Berg and Coop, 2015]. Patterns of diversity surrounding the selected site begin to resemble those expected under neutrality as the number of unique haplotypic backgrounds carrying the beneficial allele (the softness of the sweep) increases, potentially obscuring the presence of the sweep. Accordingly, the effect of a soft sweep may be unnoticeable, even if the selected allele has reached fixation.

Popular modern methods for identifying recent selective sweeps from haplotype data identify distortions in the haplotype structure following a sweep, making use of either the signature of elevated LD or reduced haplotypic diversity surrounding the site of selection. Methods in the former category [Kelly, 1997, Kim and Nielsen, 2004, Pavlidis et al., 2010] can detect both hard and soft sweeps, as neighboring neutral variants hitchhike to high frequency under either scenario. Indeed, LD-based methods may have an increased sensitivity to softer sweeps [Pennings and Hermisson, 2006b], especially relative to methods that do not use haplotype data, such as composite likelihood approaches [Kim and Stephan, 2002, Nielsen et al., 2005, Chen et al., 2010, Vy and Kim, 2015, Racimo, 2016]. Haplotype homozygosity-based methods include iHS [Voight et al., 2006], its extension, *n*S_*L*_ [Ferrer-Admetlla et al., 2014], and H-scan [Schlamp et al., 2016]. These approaches identify a site under selection from the presence of a high-frequency haplotype. Additionally, Chen et al. [2015] developed a hidden Markov model-based approach that similarly identifies sites under selection from the surrounding long, high-frequency haplotype.

While the aforementioned methods are powerful tools for identifying selective sweeps in the genome, they lack the ability to distinguish between hard and soft sweeps. It is this concern that Garud et al. [2015] address with the statistics H12 and H2/H1. H12, a haplotype homozygosity-based method, identifies selective sweeps from elevated expected haplotype homozygosity surrounding the selected site. It is computed as the expected haplotype homozygosity, with the frequencies of the two most frequent haplotypes pooled into a single frequency:

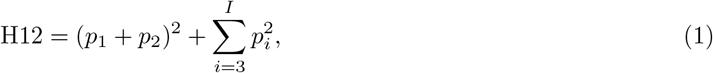

where there are *I* distinct haplotypes in the population, and *p_i_* is the frequency of the *i*th most frequent haplotype, with *p*_1_ ≥ *p*_2_ ≥ ⋯ ≥ *p_I_*. Pooling the two largest haplotype frequencies provides little additional power to detect hard sweeps relative to H1, the standard measure of expected haplotype homozygosity, where 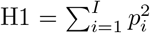 (Figure 1A, left panel). However, pooling provides more power to detect soft sweeps, in which at least two haplotypes rise to high frequency, and the distortion of their joint frequency produces an elevated expected haplotype homozygosity consistent with a sweep (Figure 1A, right panel).

**Figure 1:**
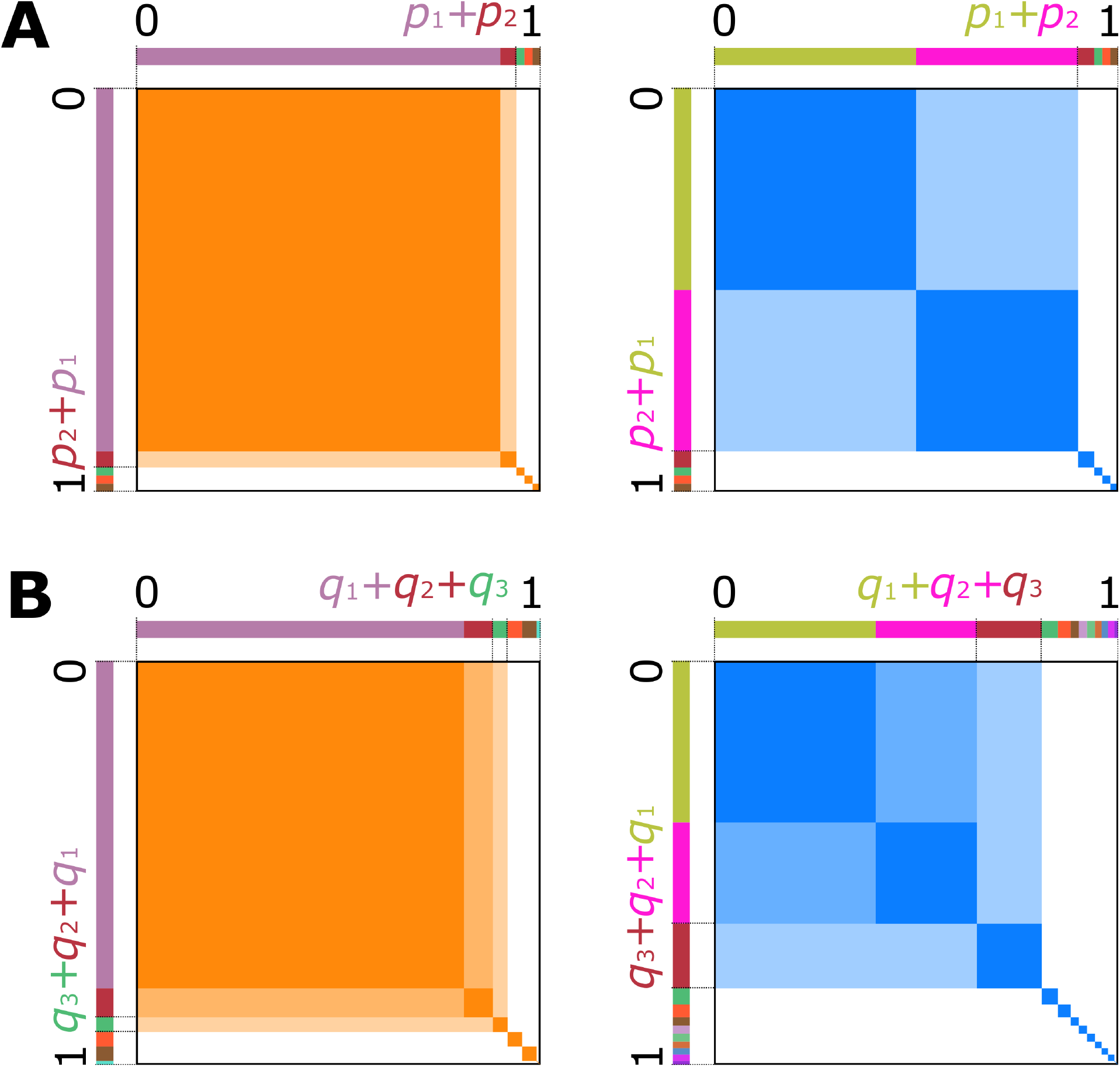
Visual representation of expected homozygosity statistics. For all panels, total area of the orange or blue squares within a panel represents the value of expected homozygosity statistics. Hard sweep scenarios are in orange, and soft sweeps are in blue. (A) Under a hard sweep (left), a single haplotype rises to high frequency, *p*_1_, so the probability of sampling two copies of that haplotype is *p*^2^. Choosing *p*_1_ as the largest frequency yields H1 (dark orange area), while pooling *p*_1_ + *p*_2_ as the largest frequency yields H12 (total orange area). Under a soft sweep (right), pooling the largest haplotype frequencies results in a large shaded area, and therefore H12 has a similar value for both hard and soft sweeps. (B) Under Hardy Weinberg equilibrium, a single high-frequency haplotype produces a single high-frequency MLG (frequency *q*_1_). Pooling frequencies up to *q*_3_ has little effect on the value of the statistic, thus G1, G12, and G123 have similar values. When two haplotypes exist at high frequency, three MLGs exist at high frequency. Under a soft sweep, pooling the largest two MLGs (G12) may provide greater resolution of soft sweeps than not pooling (G1), and pooling the largest three creates a statistic (G123) truly analogous to H12.

In conjunction with an elevated value of H12, the ratio H2/H1 serves as a measure of sweep softness, and is not meaningful on its own. H2 is the expected haplotype homozygosity, omitting the most frequent haplotype, computed as 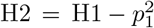, and is larger for softer sweeps. In the case of a soft sweep, the frequencies of the first- and second-most frequent haplotypes are both large, and omitting the most frequent haplotype still yields a frequency distribution in which one haplotype predominates. Under a hard sweep, the second through *I*th haplotypes are likely to be at low frequency and closer in value, such that their expected homozygosity is small. Thus, while H2 < H1 in all cases, the value of H2 is closer to that of H1 under a soft sweep.

To leverage the power of H12 and H2/H1 to detect sweeps in nonmodel organisms, for which phased haplotype data are often unavailable, we extend the application of these statistics to unphased multilocus genotype (MLG) data as G12 and G2/G1. MLGs are single strings representing a diploid individual’s allelic state at each site as homozygous for the reference allele, homozygous for the alternate allele, or heterozygous. Similarly to H12, we define G12 as

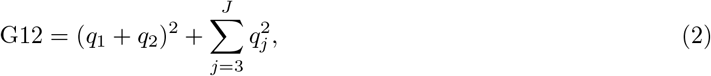

where there are *J* distinct unphased MLGs in the population, and *q_j_* is the frequency of the *j*th most frequent MLG, with *q*_1_ ≥ *q*_2_ ≥ ⋯ ≥ *q_J_*. As with haplotypes, pooling the most frequent MLGs only provides marginally more resolution to detect hard sweeps, as only a single predominant unphased MLG is expected under random mating (Figure 1B, left panel). However, because the input data for G12 and G2/G1 are unphased MLGs, we define another statistic that is uniquely meaningful in this context. The presence of multiple unique frequent haplotypes under a soft sweep implies not only that the frequency of individuals homozygous for these haplotypes will be elevated, but also that the frequencies of their heterozygotes will be elevated. When haplotypes *X* and *Y* both exist at high frequency, diploid individuals of type *XX*, *YY*, and *XY* will also exist at high frequency, assuming individuals mate randomly within the population (Figure 1B, right panel). Therefore, we can define a statistic truly analogous to H12 for unphased MLG data, G123. This statistic is calculated as

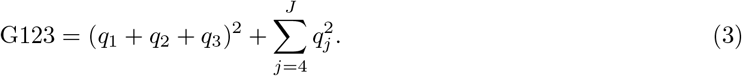

We note, however, that with this approach we do not explicitly enforce a constraint on the presence of particular high-frequency MLGs in the sample. That is, we only assume that the presence of one or more high-frequency MLGs implies a sweep, even if any one of the *XX*, *YY*, or *XY* MLGs is absent.

We show through simulation and empirical application that the statistics G12 and G123, in conjunction with the ratio G2/G1, both maintain the power of H12 to detect and classify sweeps, without requiring phased haplotype input data. Furthermore, as a closer analogue to H12, the use of G123 with G2/G1 more closely maintains the classification ability of H12 with H2/H1 than does G12. Generally, we find that the selective events visible with H12 in phased haplotype data are visible to G12 and G123 in unphased MLG data, with trends in power and genomic signature of the applications remaining consistent with one another. Accordingly, we recover well-documented sweep signatures at *LCT* and *SLC24A5* in individuals with European ancestry [Bersaglieri et al., 2004, Sabeti et al., 2007, Gerbault et al., 2009], with the latter also detected in South Asian individuals [Coop et al., 2009, Mallick et al., 2013], as well as the region linked to *EDAR* in East Asian populations [Fujimoto et al., 2007, Bryk et al., 2008, Pickrell et al., 2009], and *SYT1* in African individuals [Voight et al., 2006]. In addition, we identify novel candidates *RGS18* in African individuals, *P4HA1* in South Asian individuals, and *FMNL3* in East Asian individuals.

## Results

To detect selective sweeps, we must have power to identify loci with elevated haplotype homozygosity relative to expectations under neutral demographic scenarios. We compared the power of the MLG-based methods G12 and G123 to that of the haplotype-based methods H12 and H123 [Garud et al., 2015], at the 1% false positive rate (FPR) obtained from simulations under neutral demographic models (see *Materials and methods*). We performed simulations under population-genetic parameters inferred for human data [Takahata et al., 1995, Nachman and Crowell, 2000, Payseur and Nachman, 2000] with the forward-time simulator SLiM 2 [Haller and Messer, 2017]. Because SLiM outputs paired phased haplotypes for each diploid individual, we manually merged each individual’s haplotypes to apply the MLG-based methods. Our simulated replicates included scenarios of selective neutrality, hard sweeps, and soft sweeps. We evaluated methods across simulations of constant demographic history, as well as realistic human models of bottleneck and expansion [Lohmueller et al., 2009] (Figure 2). We then use an approximate Bayesian computation (ABC) approach to evaluate the ability of the MLG-based methods with G2/G1, and the haplotype-based methods with H2/H1, to differentiate between hard and soft sweeps. Finally, we evaluated empirical data from the 1000 Genomes Project [Auton et al., 2015], manually merging each study individual’s phased haplotypes into MLGs to observe the effect of phasing on our ability to detect selective events. See *Materials and methods* for a detailed explanation of experiments.

**Figure 2:**
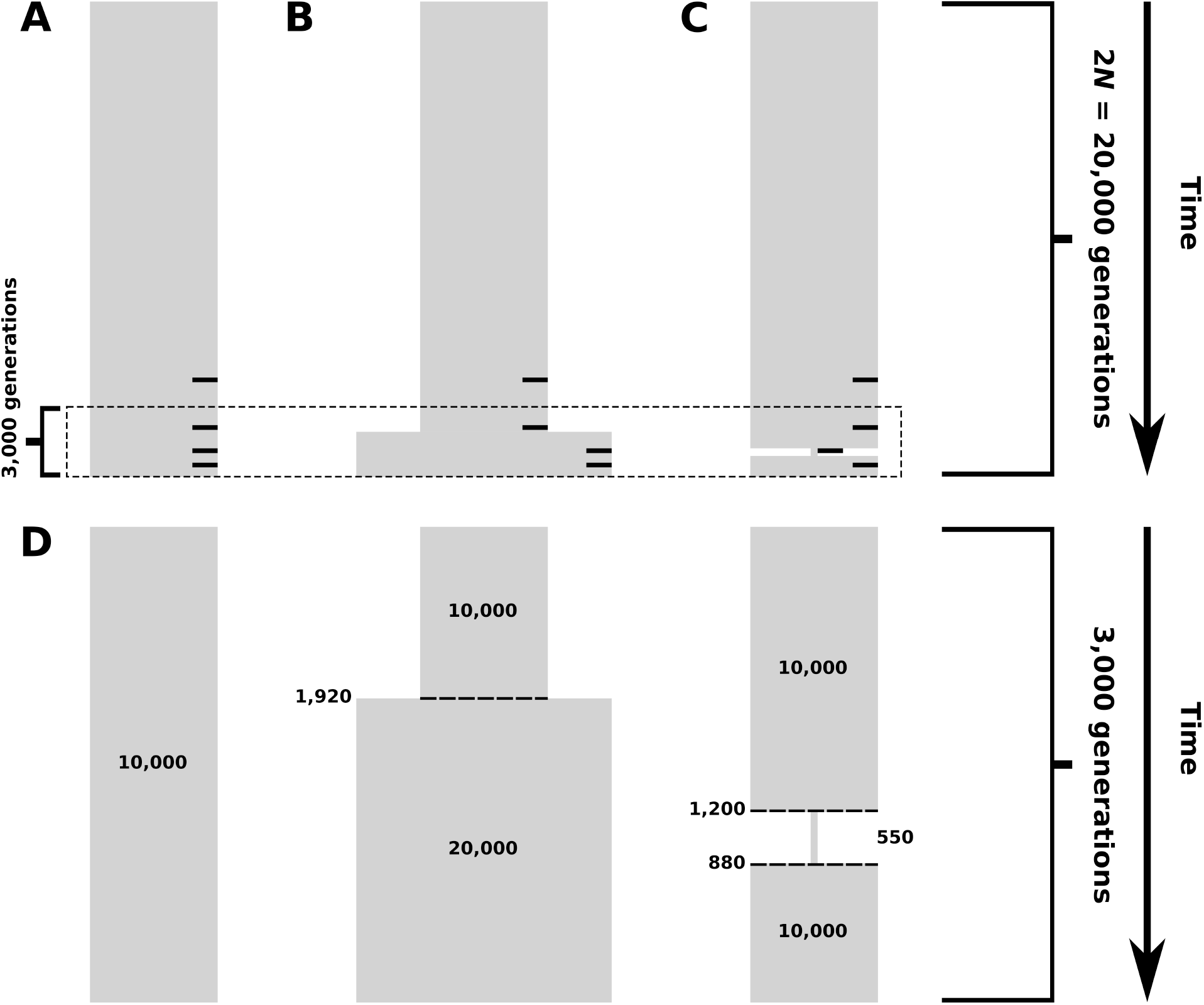
Simulated demographic models. Selection events, where applicable, occurred within 2N generations of sampling, indicated by small black bars on the right side of panels A-C corresponding to selection 4,000, 2,000, 1,000, and 400 generations before sampling. (A) Constant-size model. Diploid population size is 10^4^ individuals throughout the time of simulation. (B) Model of recent population expansion. Diploid population size starts at 10^4^ individuals and doubles to 2 × 10^4^ individuals 1,920 generations ago. (C) Model of a recent strong population bottleneck. Diploid population size starts at 10^4^ individuals and contracts to 550 individuals 1,200 generations ago, and subsequently expands 880 generations ago to 10^4^ individuals. (D) View of the final 3,000 generations across demographic models, highlighting the effects of changing demographic factors on simulated populations.

### Using G12 and G123 to detect sweeps

We demonstrate the range of sensitivity of G12 and G123 relative to H12 and H123 for selective sweeps occurring at time points between 400 and 4,000 generations before the time of sampling. We evaluated G123 to determine whether it is a more direct analogue of H12 as we expected, while our application of H123 follows from the work of Garud et al. [2015], which suggested that H123 yields little difference in power to detect sweeps relative to H12 for given sample and window size parameters. In the following experiments, we simulated 100 kilobase (kb) chromosomes carrying a selected allele at their center (sweep simulations), or carrying no selected allele for neutrality, performing 10^3^ replicates for each scenario with sample size *n* = 100 diploid individuals.

For each series of simulations, we detected sweeps using a sliding window of size 40 kb shifting by 4 kb increments across the chromosome. We selected this window size to ensure that the effect of short-range LD would not inflate the values of our statistics (Figure S1). This additionally matched the window size we selected for analysis of empirical data in non-African populations (see *Analysis of empirical data for signatures of sweeps*). According to theoretical expectations [Gillespie, 2004, Garud et al., 2015, Hermisson and Pennings, 2017], a window of size 40 kb under our simulated parameters is sensitive to sweeps with selection strength *s* ≥ 0.004 (see *Materials and methods*). Additionally, although we used a nucleotide-delimited window in our analysis, one can also fix the number of single-nucleotide polymorphisms (SNPs) included in each window (SNP-delimited window), though this somewhat changes the properties of the methods (see *Discussion*). A SNP-delimited window corresponding to approximately 40 kb for our simulated data contains on average 235 SNPs under neutrality. To supplement experiments measuring the power of each method, we also assessed the genomic distribution of G12 and G123 values to characterize their patterns under sweep scenarios.

### Tests for detection of hard sweeps

Methods that detect selective sweeps typically focus on the signature of hard sweeps, though many can detect soft sweeps as well. Accordingly, we began by measuring the ability of G12, G123, H12, and H123 to detect both partial and complete hard sweeps, under scenarios in which a single haplotype acquires a selected mutation and rises in frequency. We examined selection start times (*t*) of 400, 1,000, 2,000, and 4,0 generations before the time of sampling. These values of *t* span the time periods of various sweeps in human history [Przeworski, 2002, Sabeti et al., 2007, Beleza et al., 2012, Jones et al., 2013, Clemente et al., 2014, Fagny et al., 2014]. For each *t*, we simulated hard sweeps under the aforementioned parameters to sweep frequencies (*f*) between 0.1 and 1 for the selected allele (Figures 3 and S2). Sweeps to smaller *f* have a smaller effect on the surrounding expected haplotype homozygosity and are more difficult to detect. We performed hard sweep simulations for a large selection coefficient of *s* = 0.1 and a more moderate selection coefficient of *s =* 0.01.

**Figure 3:**
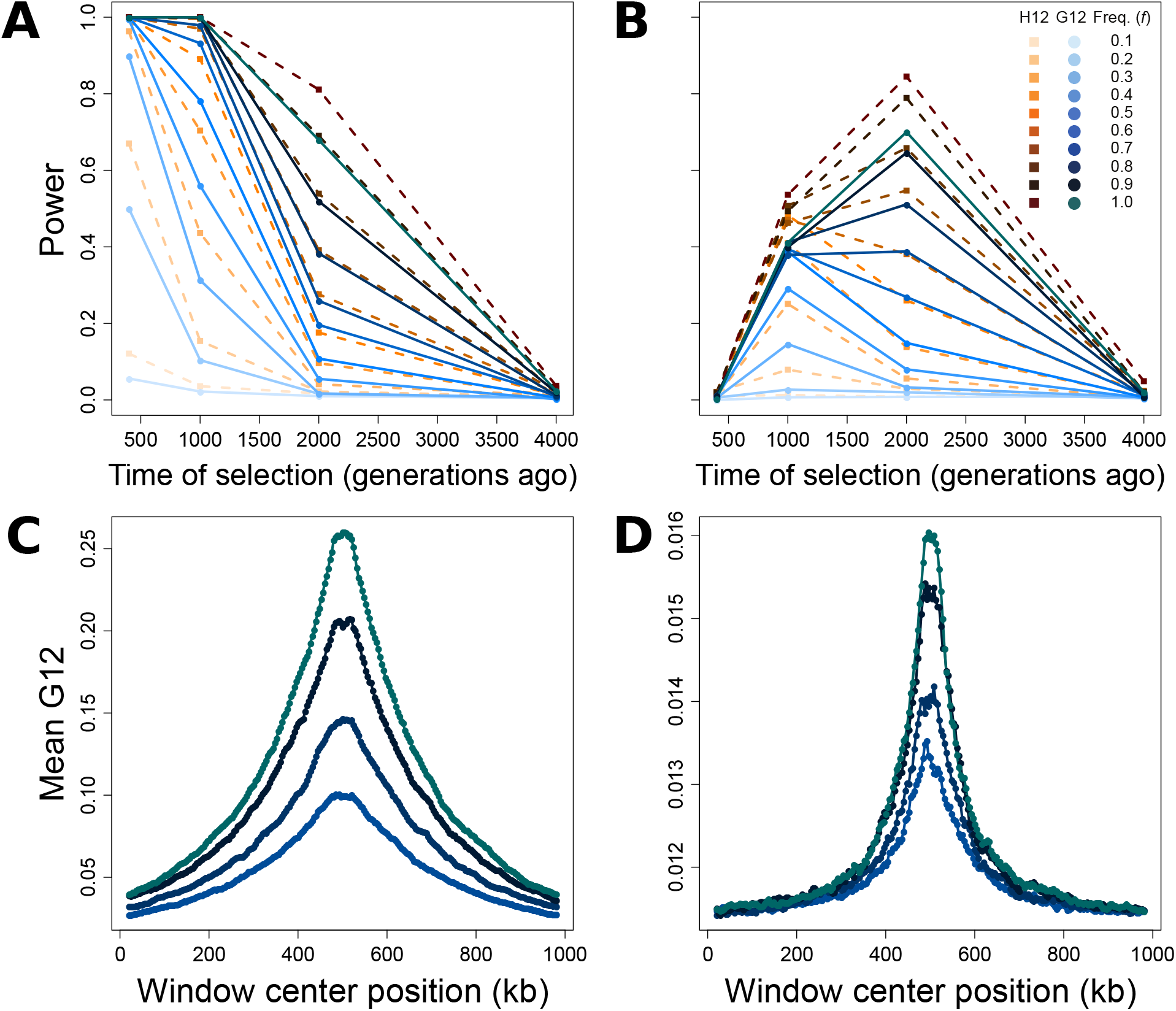
Capabilities of H12 (orange) and G12 (blue) to detect hard sweeps from simulated chromosomes, sample size *n* = 100 diploids, and window size of 40 kb for selection across four time points (400, 1,000, 2,000, and 4,000 generations before sampling) and 10 sweep frequencies (*f*, frequency to which the selected allele rises before becoming selectively neutral). Selection simulations conditioned on the beneficial allele not being lost. (A) Powers at a 1% false positive rate (FPR) of H12 and G12 to detect strong sweeps (*s* = 0.1) in a 100 kb chromosome. (B) Powers at a 1% FPR of H12 and G12 to detect moderate sweeps (*s* = 0.01) in a 100 kb chromosome. (C) Spatial G12 signal across a one Mb chromosome for strong sweeps occurring 400 generations prior to sampling. (D) Spatial G12 signal across a one Mb chromosome for moderate sweeps occurring 2,000 generations prior to sampling. Lines in (C) and (D) are mean values generated from the same set of simulations as panels A and B, and contain only results for *f* ≥ 0.7. Note that vertical axes in panels C and D differ.

The values of *t* and *f* both impact the ability of methods to identify hard sweeps (Figure 3). At the 1% FPR, all methods are suited to the detection of more recent sweeps for simulated data, losing considerable power to resolve hard sweep events occurring prior to 2,000 generations before sampling, and losing power entirely for hard sweeps occuring prior to 4,000 generations before sampling. For selection within 2,000 generations of sampling, trends in the power of the MLG-based methods resemble those of the haplotype-based methods, with the power of the MLG-based methods either matching or approaching that of the haplotype-based methods for *s* = 0.1 (Figures 3A and S2A), and following similar trends in power for *s* = 0.01 (though with slightly reduced power overall; Figures 3B and S2B), indicating that the two highest-frequency MLGs and the two highest-frequency haplotypes have a similar ability to convey the signature of a sweep.

For data simulated under strong selection, *s* = 0.1 (Figure 3A), G12 and H12 achieve their maximum power for recent selective sweeps originating within the past 1,000 generations (with little to no power lost over this interval for sweeps to large *f*). This result is expected because sweeps with such a high selection coefficient quickly reach fixation, at which point mutation and recombination break down tracts of elevated expected homozygosity until the signal fully decays, obscuring more ancient events. For a given value of *s*, selective sweeps to larger values of *f* for the selected allele additionally produce a stronger signal because more diversity is ablated the longer a sweep lasts. Thus, G12 and H12 are best able to detect sweeps over recent time intervals, especially as the sweep goes to larger values of *f*. Strong hard sweeps additionally create a peak in signal surrounding the site of selection that increases in magnitude with increasing duration of a sweep. This signal is broad and extends across the one Mb interval that we modeled in Figure 3C. These patterns repeat for G123 and H123 (Figure S2A), yielding little difference in power between H12 and H123, and no difference in power between G123 and G12 (along with a nearly-identical spatial signature along the chromosome; Figure S2C).

At a smaller selection coefficient of *s* = 0.01 (Figure 3B), G12 and H12 have a distinct range of sweep detection from *s* = 0.1. The reduced strength of selection here leads beneficial mutations to rise more slowly in frequency than for stronger selection. Consequently, after 400 generations of selection, the distribution of haplotype (and therefore MLG) frequencies has scarcely changed from neutrality, and G12 and H12 cannot reliably detect the signal of a sweep. However, the powers of G12 and H12, as well as G123 and H123 (Figure S2B), are greatest for a moderate sweep to *f* ≥ 0.9 starting 2,000 generations prior to sampling. As with stronger selection, pooling the three largest frequencies had little effect on power relative to pooling the two largest frequencies (Figure S2). We could not detect adaptive mutations appearing more anciently than 2,0 generations before sampling, indicating that all methods lose power to detect sweeps for smaller values of *s*, and that haplotype methods may outperform MLG methods for smaller values of *s* as well. Furthermore, the range of time over which methods detect a sweep narrows and shifts to more ancient time periods with decreasing *s*. Weaker selection nonetheless produces a signal peak distinct from the neutral background and proportional in magnitude to the value of *f* (Figures 3D and S2D), though expected haplotype homozygosity, and therefore expected MLG homozygosity, is reduced for moderate selection (compare vertical axes of Figures 3C and D and of Figures S2C and D).

### Tests for detection of sweeps on standing variation

We characterized the properties of G12, G123, H12, and H123 for simulated soft sweeps from selection on standing genetic variation (SSV). We generated results analogous to those for hard sweeps: measures of power for each method, and the chromosome-wide spatial distribution of the G12 and G123 signals. Across identical times of selection (*t*) and selection coefficients (*s*) as for hard sweep simulations, we simulated SSV scenarios by introducing the selected mutation on multiple haplotypes simultaneously. We evaluated method ability to correctly distinguish sweeps on *k* = 2, 4, 8, 16, and 32 initially-selected different haplotypes from neutrality. One copy of the selected allele is guaranteed to remain in the population for the entire simulation, but we do not condition on the number of sweeping haplotypes at the time of sampling. Indeed, we do not expect that for larger values of *k*, all haplotypes carrying the selected allele will remain at high frequency, or remain at all by the time of sampling (Figure S4). For our scaled (see *Materials and methods*) simulated population size of 500 diploids (unscaled 10^4^ diploids), this corresponds to having the beneficial allele present on 0.2 to 3.2% of haplotypes at the onset of selection. Our results for these tests mirror those for hard sweeps, with stronger selection on fewer distinct haplotypes yielding the most readily detectable genomic signatures (Figures 4 and S3).

**Figure 4:**
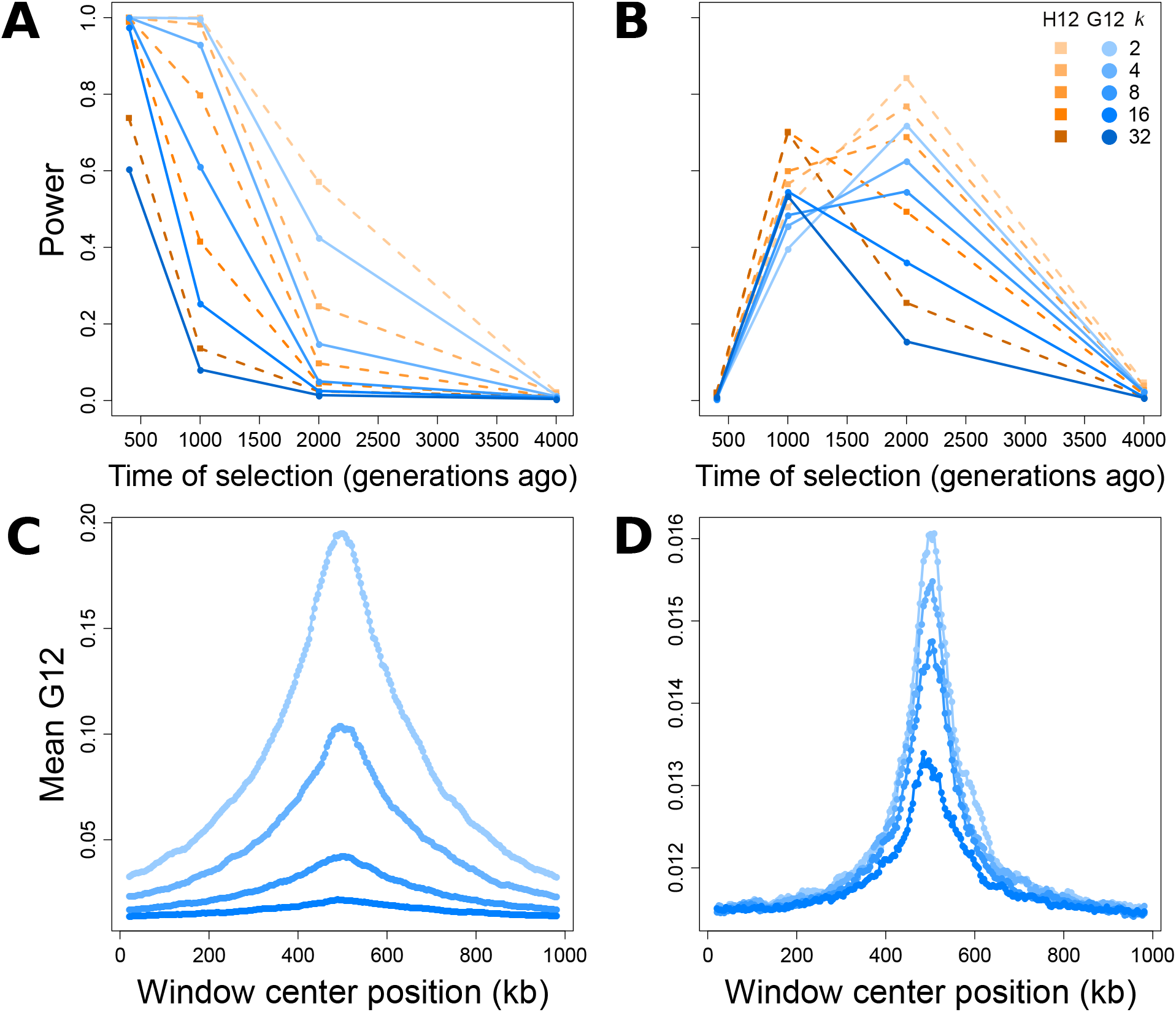
Capabilities of H12 (orange) and G12 (blue) to detect soft sweeps (SSV) from simulated chromosomes generated for selection times, sample size, and window size as in Figure 3, and five initially-selected haplotype values (*k*, number of haplotypes on which the selected allele arises at time of selection). Selection simulations conditioned on the beneficial allele not being lost. (A) Powers at a 1% false positive rate (FPR) of H12 and G12 to detect strong sweeps (*s* = 0.1) in a 100 kb chromosome. (B) Powers at a 1% FPR of H12 and G12 to detect moderate sweeps (*s* = 0.01) in a 100 kb chromosome. (C) Spatial G12 signal across a one Mb chromosome for strong sweeps occurring 400 generations prior to sampling. (D) Spatial G12 signal across a one Mb chromosome for moderate sweeps occurring 2,000 generations prior to sampling. Lines in (C) and (D) are mean values generated from the same set of simulations as panels A and B, and contain only results for *k* ≤ 16. Note that vertical axes in panels C and D differ.

SSV once again produces a signal of elevated MLG homozygosity for *s* = 0.1 that all methods most readily detect if it is recent, and rapidly lose power to detect as *t* increases. G12 and H12 reliably detect signals of SSV in simulated 100 kb chromosomes, retaining power for SSV on as many as *k* ≤ 16 haplotypes within the first 400 generations after the start of selection (Figure 4A). However, the relatively smaller expected homozygosity under SSV leads the power of each method to decay more rapidly than under a hard sweep. The levels of expected homozygosity produced under SSV are consequently smaller in magnitude than those generated under hard sweeps, but unambiguously distinct from neutrality for at least one combination of each tested *k* and *t*, with *k* = 2 most closely resembling a hard sweep throughout (Figure 4C). As with the hard sweep scenario, G123 and H123 yield little change in resolution for detecting strong soft sweeps from SSV, suggesting that the third-most frequent haplotype may have little importance in detecting sweeps (Figures S3A and C). Once again, H123 maintains slightly greater power than does G123.

G12 and H12 perform comparably well for moderate (*s* = 0.01) sweeps from SSV (Figure 4B). Similarly to hard sweep scenarios for *s* = 0.01, G12 and H12 detected soft sweeps from SSV occurring between 1,000 and 2,000 generations before sampling. Once again, the power of H12 was greater than that of G12, with trends in power for G12 following those of H12. For both MLG and haplotype data, the inclusion of additional selected haplotypes at the start of selection up to *k* = 8 only slightly reduced the maximum power of G12 and H12 to detect sweeps, but with time at which maximum power is reached shifting from 2,000 generations before sampling for *k* ≤ 8 to 1,000 generations before sampling for *k* ≥ 16. Additionally, the spatial signal for moderate sweeps was comparable between SSV and hard sweep scenarios (Figure 4D). This result may be because at lower selection strengths, haplotypes harboring adaptive alleles are more likely to be lost by drift, leaving fewer distinct selected haplotypes rising to appreciable frequency. These trends persist for G123 and H123, which display similar powers to G12 and H12 across all scenarios (Figures S3B and D).

### Effect of population size changes on detection capabilities of G12 and G123

Changes in population size that occur simultaneously with or after the time of selection may impact the ability of methods to detect sweeps because haplotypic diversity may decrease under a population bottleneck, or increase under a population expansion [Campbell and Tishkoff, 2008]. To test the robustness of the expected homozygosity statistics to these potentially confounding scenarios, we modeled hard sweeps following the human population bottleneck and expansion parameters inferred by Lohmueller et al. [2009] (Figure 2). We measured the powers of the MLG- and haplotype-based methods across our previously-tested parameters, using simulated 100 kb chromosomes and sliding windows, and approaching these scenarios in two ways.

First, we applied a 40 kb window as previously to evaluate the effect of population size change on the power of expected homozygosity methods. Under a bottleneck, a 40 kb window is expected to carry fewer SNPs than under a constant-size demographic history, whereas an expansion results in greater diversity per window. Second, we examined whether we could increase the robustness of the expected homozygosity methods to population size changes by adjusting the window size for each scenario to match the expected number of segregating sites for a 40 kb window under constant demographic history. To do this, we followed the approach outlined in DeGiorgio et al. [2014], increasing window size for bottleneck simulations and decreasing window size for expansion simulations. We employed windows of size 56,060 nucleotides for bottleneck, and of size 35,048 nucleotides for expansion scenarios [see DeGiorgio et al., 2014].

A recent population bottleneck reduces the powers of all methods to detect sweeps, whereas a recent population expansion enhances power (Figures S5 and S6). This results from the genome-wide reduction in haplotypic diversity under a bottleneck relative to the constant-size demographic history. Thus, the maximum values of the expected homozygosity statistics in the absence of a sweep are inflated, resulting in a distribution of maximum values under neutrality that has increased overlap with the distribution under selective sweeps. In contrast, haplotypic diversity is greater under the population expansion than what is expected for the constant-size demographic history, rendering easier the detection of elevated expected homozygosity due to a sweep.

For strong selection (*s* = 0.1) under a population bottleneck, all methods using unadjusted windows have reliable power to detect only recent hard sweeps to large *f* occurring within 1,000 generations of sampling (Figures S5A and S6A). Adjusting window size has little effect on this trend, with powers for sweeps beginning 400 generations before sampling increasing only slightly (Figures S5C and S6C). This result indicates that we can apply the expected homozygosity methods to populations that have experienced a severe bottleneck and make accurate inferences about their selective histories. Similarly, adjusting window size had little effect on the power of methods to detect a sweep under a population expansion, wherein power is already elevated. As with the bottleneck scenario, reducing the size of a 40 kb window (Figure S5B and S6B) to 35,048 bases (Figure S5D and S6D) provided a minor increase in power to detect selective events occurring within 2,000 generations of sampling, with high power for larger values of *f* extending to 2,000 generations prior to sampling.

### Distinguishing hard and soft sweeps with G2/G1

Having identified selective sweeps with the statistics G12 or G123, our goal is to make an inference about the number of sweeping haplotypes. To distinguish between hard and soft sweeps, Garud et al. [2015] defined the ratio H2/H1, which is larger under a soft sweep and smaller under a hard sweep. The H2/H1 ratio leverages the observation that haplotypic diversity following a soft sweep is greater than that under a hard sweep. Garud and Rosenberg [2015] showed that the value of H2/H1 is inversely correlated with that of H12, and that identical values of H2/H1 have different interpretations depending on their associated H12 value. Therefore, H2/H1 should only be applied in conjunction with H12 when H12 is large enough to be distinguished from neutrality.

Here, we extend the application of H2/H1 to MLGs. As with the haplotype approach, G2/G1 is larger under a soft sweep and smaller under a hard sweep, because MLG diversity following a soft sweep is greater than under a hard sweep. G2/G1 should therefore distinguish between hard and soft sweeps similarly to H2/H1, conditional on a high G12 or G123 value. To demonstrate the classification ability of the MLG-based methods with respect to the haplotype-based methods, we began by generating 10^6^ simulated replicates of 40 kb chromosomes with sample size *n* = 100 diploids for hard sweep and SSV scenarios, treating each chromosome as a single window and recording its G12, G123, and G2/G1 values (see *Materials and methods*).

We evaluated the ability of G2/G1 with G12 or G123 to distinguish between hard sweeps and soft sweeps from SSV specifically from *k* = 3 and *k* = 5 drawn haplotypes, both within the range of method detection (Figures 4 and S3), with all sweeps allowed but not guaranteed to go to fixation. We examined two values of *k*, distinct from one another and from hard sweeps, to illustrate the effect of model choice on sweep classification. Each experiment evaluated the likelihood that a soft sweep scenario would produce a particular paired (G12, G2/G1) or (G123, G2/G1) value relative to a hard sweep scenario. We measured this relative likelihood by plotting the Bayes factors (BFs) for paired (G12, G2/G1) and (G123, G2/G1) test points generated from an approximate Bayesian computation (ABC) approach (see *Materials and methods*). A BF > 1 indicates a greater likelihood of a soft sweep generating the paired values of a test point, and a BF < 1 indicates that a hard sweep is more likely to have generated such values. In practice, however, we only assign BF ≤ 1/3 as hard and BF ≥ 3 as soft to avoid making inferences about borderline cases (Figure 5). For each replicate, time of selection (*t*) and selection strength (*s*) were drawn uniformly at random on a log-scale from *t* ∈ [40, 2000] generations before sampling and *s* ∈ [0.005, 0.5].

**Figure 5:**
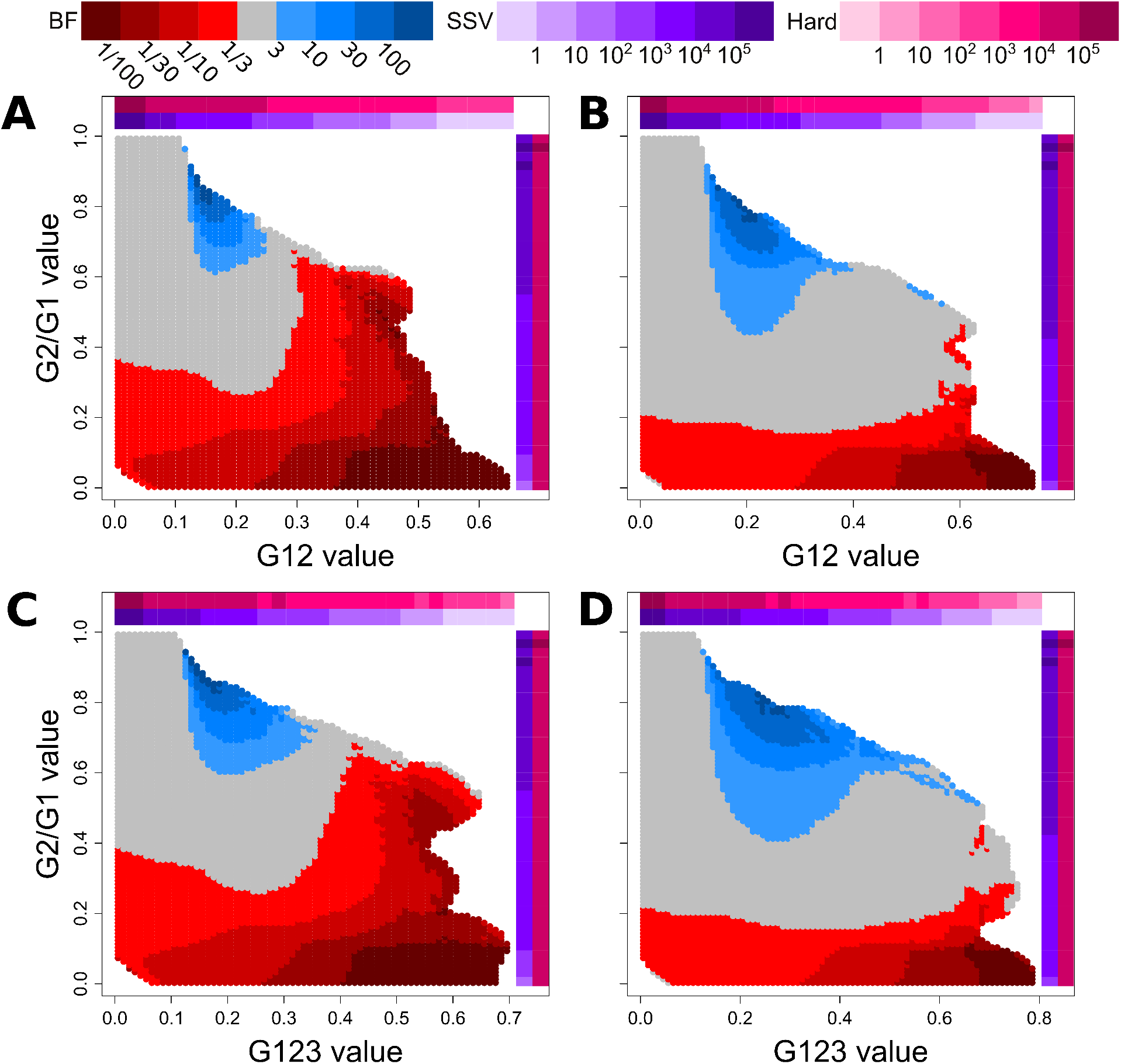
Assignment of Bayes factors (BFs) to tested paired values of (G12, G2/G1) and (G123, G2/G1). Plots represent the relative probability of obtaining a paired (G12, G2/G1) or (G123, G2/G1) value within a Euclidean distance of 0.1 from a test point for hard versus soft sweeps, determined as described in the *Materials and methods*. Selection coefficients (*s*) and times (*t*) were drawn as described in the *Materials and methods*. Red regions represent a higher likelihood for hard sweeps, while blue regions represent a higher likelihood for soft sweeps. Colored bars along the axes indicate the density of G12 or G123 (horizontal) and G2/G1 (vertical) observations within consecutive intervals of size 0.025 for hard sweep (magenta) and SSV (purple) simulations. (A) BFs of paired (G12, G2/G1) values for hard sweep scenarios and SSV scenarios (*k* = 5). (B) BFs of paired (G12, G2/G1) values for hard sweep scenarios and SSV scenarios (*k* = 3). (C) BFs of paired (G123, G2/G1) values for hard sweep scenarios and SSV scenarios (*k* = 5). (D) BFs of paired (G123, G2/G1) values for hard sweep scenarios and SSV scenarios (*k* = 3). Only test points for which at least one simulation of each type was within a Euclidean distance of 0.1 were counted (and therefore colored).

The comparison of hard sweep and SSV scenarios provides a distribution of BFs broadly in agreement with expectations for the haplotype-based approaches (Garud et al. [2015], Garud and Rosenberg [2015]; Figure 5). In Figure 5, colored in blue are the values most likely to be generated under SSV, and colored in red are the values most likely to be generated under hard sweeps. In all scenarios tested, hard sweeps produce relatively smaller G2/G1 values than do soft sweeps. Intermediate G12 and G123 paired with large values of G2/G1 are more likely to result from soft sweeps than from hard sweeps. SSV cannot generate large values of G12 or G123 because these sweeps are too soft to elevate homozygosity levels to the extent observed under hard sweeps. This is particularly so when soft sweeps are simulated with *k* = 5. Therefore, the majority of test points with extreme values of G12 and G123, regardless of G2/G1, have BF ≤ 1/3 (meaning only one SSV observation within a Euclidean distance of 0.1 for every three or more hard sweep observations), and this is in line with the results from the constant-size demographic model of Garud et al. [2015] for comparisons between hard sweeps and the softest soft sweeps. Additionally, we cannot classify sweeps if the values of G12 and G123 are too low, as these values are unlikely to be distinct from neutrality. Thus, our ability to distinguish between hard and soft sweeps is greatest for intermediate values of G12 and G123. In practice, our empirical top sweep candidates all converge over this range of the (G12, G2/G1) and (G123, G2/G1) values (Figure 6), meaning that we can confidently classify sweeps from outlying values of G12 and G123 in our data as hard or soft.

**Figure 6:**
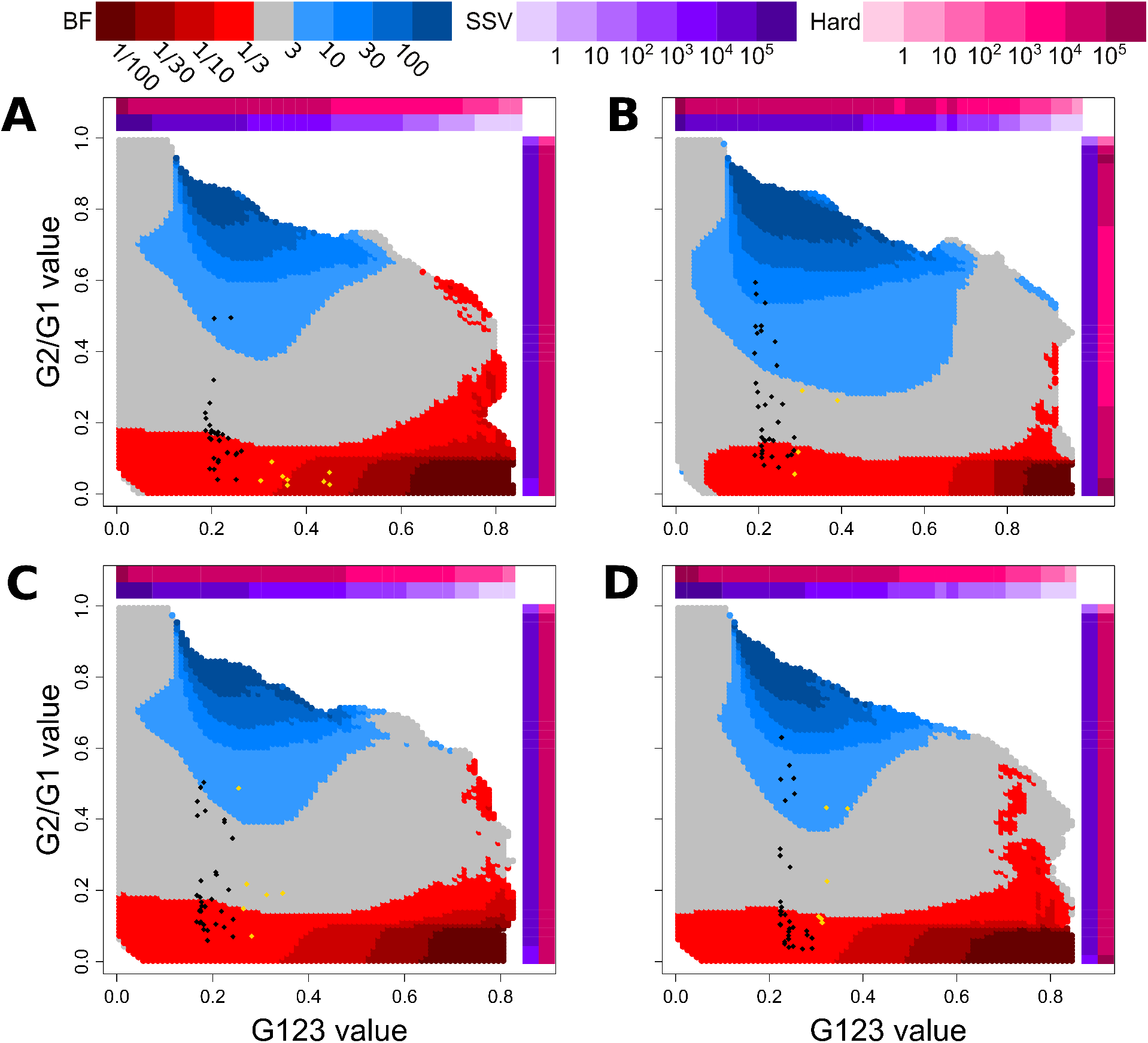
(G123, G2/G1) values used to distinguish hard (red) and soft (blue) sweeps in human empirical data using demographic models inferred with `smc++` [Terhorst et al., 2017]. Points representing the top 40 G123 selection candidates (Tables S4, S7, S10, and S13) for the (A) CEU, (B) YRI, (C) GIH, and (D) CHB populations are overlayed onto each population’s specific (G123, G2/G1) distribution. Candidates exceeding the significance threshold (Table S1; different for each population) are colored in gold. Colored bars along the horizontal (G123) and vertical (G12) axes are defined as in Figure 5.

In Figure S7, we repeat our ABC procedure for the phased haplotype data corresponding to our preceding analyses. We find that a small proportion of (G12, G2/G1) and (G123, G2/G1) values for which we lack the ability to distinguish hard and soft sweeps (gray points), corresponds to (H12, H2/H1) values that do classify sweeps as soft. Additionally, the (H123, H2/H1) values yielded a still larger proportion of SSV-classified (blue) values. This result may indicate that the haplotype approaches maintain a somewhat greater ability to classify sweeps than do the MLG approaches. Accordingly, the skew toward larger BFs among the (G123, G2/G1) values relative to (G12, G2/G1) may indicate that classification with the former may more closely resemble classification using (H12, H2/H1) values.

To further characterize the classification properties of both the MLG- and haplotype-based approaches, we next employed an alternative ABC approach in which we determined the posterior distribution of *k* for a range of (G12, G2/G1), (G123, G2/G1), (H12, H2/H1), and (H123, H2/H1) value combinations. For these experiments, we generated 5 × 10^6^ replicates of sweep scenarios with *k* ∈ {1, 2, …, 16} drawn uniformly at random for each replicate, maintaining all other relevant parameters identical to the BF experiments (see *Materials and methods*). From the posterior distribution of *k* values, we assigned the most probable *k* for a wide range of points using both MLG and haplotype data (Figure S8), and generated probability density functions across H12, H2/H1, G123, and G2/G1 for each value of *k* (Figure S9). G12, G123, H12, and H123 values were larger for sweeps with smaller *k*, and G2/G1 values were smaller for these sweeps, as expected. We achieved a finer resolution from haplotypes than from MLGs, as in the BF experiments (Figures 5 and S7), and found our inference of the most probable values of *k* across test points to be concordant with BF-based results. As previously, hard sweeps (*k* = 1) occupied larger values of G12, G123, H12, and H123 and smaller values of G2/G1 and H2/H1, with inferred *k* (similarly to inferred BF) increasing with increasing G2/G1 and H2/H1, regardless of G12, G123, H12, and H123 value. Thus, our alternative ABC approach can assign a most probable *k* from the entire tested range of *k* ∈ {1, 2, …, 16}, allowing for sweep classification without the ambiguity of BFs.

### Analysis of empirical data for signatures of sweeps

We applied G12, G123, and H12 to whole-genome variant calls on human autosomes from the 1000 Genomes Project [Auton et al., 2015] to compare the detective properties for each method on empirical data (Figures 7 and S11-S18; Tables S3-S14). This approach allowed us to understand method performance in the absence of confounding factors such as missing data and small sample size. The choice of human data additionally allowed us to validate our results from the wealth of identified candidates for selective sweeps within human populations worldwide that has emerged from more than a decade of research [e.g., Sabeti et al., 2002, Bersaglieri et al., 2004, Voight et al., 2006, Bhatia et al., 2011, Chen et al., 2015, Schrider and Kern, 2016, Cheng et al., 2017]. To apply our MLG-based methods to the empirical dataset, consisting of haplotype data, we manually merged the haplotypes for each study individual to generate MLGs. Thus, all comparisons of G12 and G123 with H12 were for the same data, as in our simulation experiments.

**Figure 7:**
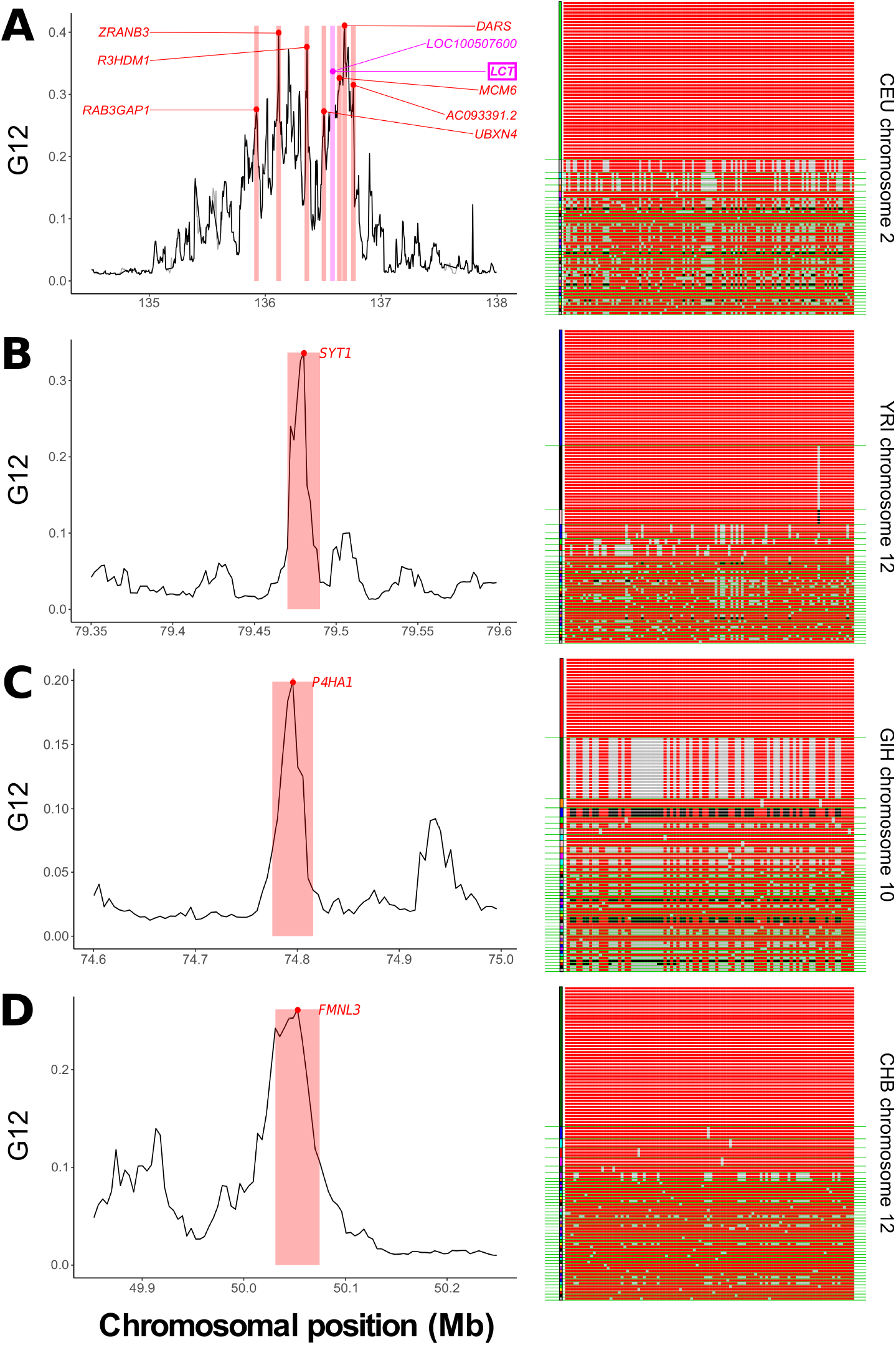
Outlying G12 signals in human genomic data. For each population, we show a top selection candidate and display its sampled MLGs within the genomic window of maximum signal. Red and black sites are homozygous genotypes at a SNP within the MLG, while gray are heterozygous. Green lines separate MLG classes in the sample. (A) CEU chromosome 2, centered around *LCT*, including other outlying loci (labeled). *LOC100507600* is nested within *LCT* (left). A single MLG exists at high frequency, consistent with a hard sweep (right). (B) YRI chromosome 12, centered on *SYT1* (left). This signal is associated with two elevated-frequency MLGs (right). (C) GIH chromosome 10, centered on *P4HA1* (left). Two MLGs exist at high frequency (right). (D) CHB chromosome 12, centered on *FMNL3* (left). A single MLG predominates in the sample (right).

For our analysis of human data, we focused on individuals from European (CEU), African (YRI), South Asian (GIH), and East Asian (CHB) descent. Across all populations, we assigned *p*-values and BFs, as well as maximum posterior estimates and Bayesian credible intervals on *k*, for the top 40 selection candidates (see *Materials and methods*). Our Bonferroni-corrected significance threshold [Neyman and Pearson, 1928] was 2.10659 × 10^−6^, with critical values for each statistic in each population displayed in Table S1. We defined soft sweeps as those with BF ≥ 3 or inferred *k* ≥ 2, and hard sweeps as those with BF ≤ 1/3 or inferred *k* = 1. Following each genome-wide scan, we filtered our raw results using a mappability and alignability measure (see *Materials and methods*), following the approach of Huber et al. [2016]. We additionally omitted genomic windows from our analysis with fewer than 40 SNPs, the expected number of SNPs in our genomic windows [Watterson, 1975] under the assumption that a strong recent sweep has affected all but one of the sampled haplotypes. This is thus a conservative approach. We display the filtered top 40 outlying sweep candidates for G12, G123, and H12, including *p*-values, BFs, and inferred *k* (with credible interval), in Tables S3-S14. We also overlay the top 40 selection candidates for each population onto (G123, G2/G1) test points (Figures 6 and S10). For all populations, we see that top candidates, regardless of assignment as hard or soft, generate broadly similar G123 values within a narrow band of paired (G123, G2/G1) values. Finally, we indicate the top 10 selection candidates in chromosome-wide Manhattan plots for both G12 and G123 (Figures S11-S18). Expectedly, G12 and G123 plots are nearly identical in their profiles.

We recovered significant signals from the well-documented region of CEU chromosome 2 harboring the *LCT* gene, which confers lactase persistence beyond childhood [Bersaglieri et al., 2004]. Although filtering removed *SLC24A5*, another expected top candidate controlling skin pigmentation, the adjacent *SLC12A1* gene remained. Assigned BFs and inferred values of *k* suggest that hard sweeps in each of these regions yield the observed signals (Tables S3 and S4). In YRI (Tables S6 and S7), we most notably found the previously-identified *SYT1, HEMGN*, and *NNT* [Voight et al., 2006, Pickrell et al., 2009, Fagny et al., 2014, Pierron et al., 2014]. *SYT1* and *HEMGN* were significant for G12, G123, and H12 analyses, with *SYT1* yielding the strongest signal by a large margin, while *NNT* was not significant. Of these, we could only confidently classify *HEMGN*, which we uniformly identified as hard. Though we were more likely to confidently classify candidate sweeps in YRI as hard from their MLG-based BFs, the proportion of top candidates assigned as hard from the posterior distribution of *k* remained comparable across data types, and generally greater than the levels we observed in other populations (see *Discussion* for further analysis). The most outlying target of selection in GIH (Tables S9 and S10) for all methods was at *SLC12A1*, a significant signal corresponding to a sweep shared among Indo-European populations [Mallick et al., 2013], which we also recovered as a top candidate in CEU. We could classify this signal as hard from haplotype data, but we assigned *k* = 2 from MLGs, despite a BF < 1. Finally, our analysis of CHB returned *EDAR-adjacent* genes among the top sweep candidates, including *LIMS1, CCDC138*, and *RANBP2* (each below the significance threshold), though not *EDAR* itself (Tables S12 and S13), and additionally *MIR548AE2* and *LONP2*, adjacent to the site of a proposed sweep on earwax texture within *ABCC11* [Ohashi et al., 2010], which we recovered as another top candidate.

In Figure 7, we highlight for each population one example of a sweep candidate, including its G12 signal profile, with the genomic window of maximum value highlighted, and a visual representation of the MLG diversity within that region. For the CEU population, we present *LCT* (*p* < 10^−6^), and additionally highlight the nearby outlying candidates, each of which was within the top 10 outlying G12 signals in the population (Figure 7A, left panel). The distribution of MLGs surrounding *LCT* in the sample showed a single predominant MLG comprising approximately half of individuals, consistent with a hard sweep (Figure 7A, right panel). Accordingly, *LCT* yielded a BF ≈ 0.1, indicating that a hard sweep is tenfold more likely to yield this signal than a soft sweep (from *k* = 5), and an inferred *k* = 1 supports this result. For the YRI population, the top selection signal for all analyses was *SYT1* (*p* = 10^−6^), previously identified by Voight et al. [2006] (Figure 7B, left panel). Here, one high-frequency and one intermediate-frequency MLG predominated in the population (Figure 7B, right panel), but we could not confidently assign the signal as hard or soft, with haplotypes suggesting *k* = 1 and MLGs suggesting *k* = 2. This is because one high-frequency haplotype exists in the population, carried by approximately half of individuals, while another haplotype exists in approximately one quarter of individuals. In GIH, we found *P4HA1* as a selection candidate exceeding the significance threshold for haplotype data (*p* = 10^−6^), but not for MLG data. Although we were unable to confidently assign the putative sweep on *P4HA1* as hard or soft from BFs, we note that two MLGs, as well as two haplotypes, exist at elevated frequency here, and that all methods yielded BF > 1 and *k* > 1, suggesting that *P4HA1* is likely the site of a soft sweep, but on fewer than *k* = 5 haplotypes (Figure 7C, right panel). Finally, our scan in CHB returned the undocumented *FMNL3* gene as a top candidate from the G12 analysis (*p* = 5 × 10^−6^; Figure 7D, left panel). A single high-frequency MLG predominated at this site, and this yielded a BF from MLG data of 0.147, and inferred *k* = 1 from all data, indicating a hard sweep (Figure 7D, right panel).

Through the application of G123 and G2/G1 we have identified and classified a number of interesting sweep candidates. We further explored the existence of a more general relationship between top sweep candidates and the prevalence and length of runs of homozygosity. Previous research has indicated that short-to-intermediate runs of homozygosity spanning tens to hundreds of kilobases are characteristic of recent sweeps [Pemberton et al., 2012, Blant et al., 2017], and we sought to examine whether there was a correlation of G123 or sweep softness (using log_10_(BF) as proxy) with the proportion of individuals falling in a run of homozygosity of specific length. To this end, we intersected our top candidates lists with the inferred coordinates of short to intermediate runs of homozygosity from Blant et al. [2017]. We found that the proportion of individuals with runs of homozygosity of intermediate length (class 4) is positively correlated (correlation coefficient = 0.32, *p*-value = 3.66 × 10^−5^) with G123 (Table S2), likely due to stronger and more recent sweeps generating larger G123. Moreover, the proportion of individuals with runs of homozygosity of intermediate length is negatively correlated (correlation coefficient = −0.26, *p*-value = 1.02 < 10^−3^) with log_10_(BF) (Table S2), likely due to the narrower genomic signature left behind by soft sweeps relative to hard sweeps. In contrast, we observe no significant correlation for smaller runs of homozygosity (classes 2 and 3), which have also been proposed to potentially be affected by selective sweeps [Pemberton et al., 2012, Blant et al., 2017].

## Discussion

Selective sweeps represent an important outcome of adaptation in natural populations, and detecting these signatures is key to understanding the history of adaptation in a population. We have extended the existing statistics H12 and H2/H1 [Garud et al., 2015] from phased haplotypes to unphased MLGs as G12, G123, and G2/G1, and demonstrated that the ability to detect and classify selective sweeps as hard or soft remains. Across simulated selective sweep scenarios covering multiple selection start times and strengths, as well as sweep types and demographic models, we found that both G12 and G123 maintain comparable power to H12. The most immediate implication of these results is that signatures of selective sweeps can be identified and classified in organisms for which genotype data are available, without the need to generate phased haplotypes. Because phasing may be difficult or impossible given the resources available to a study system, while also not being error-free [Browning and Browning, 2011, O’Connell et al., 2014, Laver et al., 2016, Castel et al., 2016, Zhang et al., 2017], the importance of our MLG-based approach is apparent. Although phased haplotypes tend to be preferable for use with expected homozygosity statistics based on our findings, we nonetheless observe a high degree of congruence in practice between the lists of selection candidates for human empirical data emerging from analyses on haplotypes and MLGs (Tables S3-S14).

### Performance of G12 and G123 for simulated data

G12 and G123, similarly to H12 and H123, are best suited to the detection of recent and strong selective sweeps in which the beneficial allele has risen to appreciable frequency. This is as expected because haplotype (and therefore MLG) homozygosity increases under sweeps, which results in a distinct signature from which to infer the sweep. This extended tract of sequence identity within the population erodes over time and returns to neutral levels due to the effects of recombination and mutation. The strength of selection and range of time over which the expected homozygosity-based methods can detect selection are inversely correlated. Our approach detects weaker selective events only if they started far enough back in time, and has a narrower time interval of detection than do stronger events (compare panels A and B across Figures 3, 4, S2, and S3). This is because alleles under weaker selection increase in frequency toward fixation more slowly than those under stronger selection, and so more time is required to generate a detectable signal. In the process, the size of the genomic tract that hitchhikes with the beneficial allele decreases due to recombination and is smaller than under a hard sweep. Panels C and D from Figures 3, 4, S2, and S3 motivate this point. Across all simulation scenarios, stronger selection produces on average a wider and larger signature surrounding the site of selection, while weaker sweeps are more difficult to detect and classify. For empirical analyses, this means we are more likely to detect stronger sweeps, as reductions in diversity from strong selection persist for hundreds of generations and can leave footprints on order of hundreds of kilobases [Gillespie, 2004, Garud et al., 2015, Hermisson and Pennings, 2017].

Expectedly, the signatures of sweeps, and the power of the expected homozygosity methods to detect them, vary across selective sweep scenarios, with nearly identical trends in haplotype and MLG data. Strong (s = 0.1) hard sweeps to high sweep frequency *f* are easiest to detect, as the single, large tract of sequence identity generated under a strong hard sweep remains distinct from neutrality for the longest time interval relative to other scenarios (Figures 3A and C and Figures S2A and C). Nonetheless, power to distinguish soft sweeps is large for the most recent simulated sweeps. Indeed, a soft sweep yields a smaller tract of sequence identity that requires a shorter time to break apart, but for strong selection on up to *k* = 16 different haplotypic backgrounds (1.6% of the total population), both the MLG and haplotype methods have perfect or nearly-perfect power (Figures 4A and S3A). While this power rapidly fades for selection within 1,000 generations of sampling for *k* > 4, our strong sweep results illustrate that selection coefficient s, more than partial sweep frequency *f* or number of initially-selected haplotypes *k*, influences the power of our pooled expected homozygosity methods, and that pooling can allow for similar detection of hard and soft sweeps. Our moderate selection (s = 0.01) results further highlight this. Once again, we see a distinct concordance in power trends between hard (Figures 3B and D and Figures S2B and D) and soft (Figures 4B and D and Figures S3B and D) sweeps that depends primarily on the value of *s* and secondarily on *f* or *k*.

Because genomic scans using G12, G123, H12 and H123 are window-based, the choice of window size is an important determinant of the methods’ sensitivity. As do Garud et al. [2015], we recommend a choice of window size that minimizes the influence of background LD on window diversity, while maximizing the proportion of sites in the window affected by the sweep. Windows that are too small may contain extended homozygous tracts not resulting from a sweep, while windows that are too large will contain an excess of neutral diversity leading to a weaker signal, while overlooking weaker selective events [Gillespie, 2004, Garud et al., 2015, Hermisson and Pennings, 2017]. Accordingly, our choice of a 40 kb sliding window to analyze simulation results derives from our observation that the value of LD between pairs of SNPs separated by 40 kb in these simulations is less than one-third of the LD between pairs separated by one kb, as measured from the squared correlation, *r*^2^ (Figure S1). We also found that for recent selection within 400 generations of sampling, power under bottleneck or expansion does not change for a 40 kb analysis window (Figures S5 and S6). This is especially important in the context of a population bottleneck, in which levels of short-range LD are elevated beyond their expected value under a constant-size demographic history [Slatkin, 2008, DeGiorgio et al., 2009]. Thus, our population size change experiments indicated that for sufficiently large analysis windows, further adjusting window size does not improve power. The trends in power that we observed for samples of *n* = 100 diploids and 40 kb genomic windows also persisted for experiments with a smaller sample size of *n* = 25 (Figure S19). The expected homozygosity methods are therefore suitable for detecting sweeps from a wide range of sample sizes, though samples need to be large enough to capture the difference in variation between selected and neutral regions of the genome, as smaller samples result in fewer sampled haplotypes [Pennings and Hermisson, 2006a]. Accordingly, the classification of sweeps requires substantially larger sample sizes, as differentiating between hard and soft sweeps requires the detection of a more subtle signal than does distinguishing selection from neutrality.

Although we exclusively used a nucleotide-delimited window in our present analyses, it is possible to search for signals of selection using a SNP-delimited window, and this was the approach of Garud et al. [2015]. Similarly to our present approach, the number of SNPs to include in a window could be determined based on the decay in pairwise LD between two sites separated by a SNP-delimited interval. Under the SNP-delimitation approach, each analyzed genomic window includes a specified number of SNPs. Thus, the range of physical window sizes may be broad. In principle, the use of a SNP-delimited window prevents the inclusion of SNP-poor windows. Accordingly, SNP delimitation may be inherently robust to the effect of bottlenecks, or to the misidentification of heterochromatic regions as sweep targets. In practice, however, we can filter out nucleotide-delimited genomic windows carrying too few SNPs to overcome confounding signals. More importantly, allowing for a variable number of SNPs in a window allows the genomic scan to identify sweeps not only from distortions in the haplotype frequency spectrum, but also from reductions in the total number of distinct haplotypes, which are more constrained in their range of values when conditioned on a specific number of SNPs. Because both of these signatures can indicate a sweep, it may be useful to consider each. Even so, the use of a SNP-delimited window may be preferable for SNP chip data. That is, SNP density can be low relative to whole-genome data, resulting in an excess of regions spuriously appearing to be under selection within a nucleotide-delimited window. Indeed, Schlamp et al. [2016] employ a SNP-delimited window approach for their canine SNP array dataset.

During a genomic scan, it may also be helpful to account for sources of uncertainty in the data. Foremost among these is uncertainty in genotype calls [Marchini and Howie, 2010, Nielsen et al., 2011]. Modern genotype calling methods provide a posterior probability for each genotype [He et al., 2014, Korneliussen et al., 2014, Fumagalli et al., 2014], and so it may be possible to assign to each analysis window a weighted mean G12 or G123 score from this posterior to produce a more accurate representation of sweep events throughout the study population’s genome. It is also possible that windows of elevated G12 and G123 value may arise in the absence of random mating. That is, although our approach assumes elevated MLG homozygosity derives from elevated haplotype homozygosity as a result of random mating, we do not specifically evaluate whether observed patterns of MLG diversity are compatible with the random mating assumption. Such an approach could condition on the presence of one high-frequency MLG with only homozygous sites in the case of a hard sweep, or at least two high-frequency homozygous MLGs in the case of a soft sweep. To further consider this point, we rescanned the 1000 Genomes dataset, but randomly paired haplotypes into diploid MLGs to simulate random mating. Our lists of outlying sweep candidates for G123 across each study population after random reshuffling were highly concordant with the lists for the true set of diploid individuals (Tables S5, S8, S11, and S14).

While power to detect hard and soft sweeps is comparable, the possible values of G12 and G2/G1 that can be generated under hard versus soft sweeps for a variety of *k* values are distinct. Thus, we can properly classify sweeps from MLG data (Figure 5, 6, S8, and S10). This result matched our theoretical expectations (Figure 1), and corresponded to the results from haplotype data as well (Figure S7). However, we note that with the BF-based ABC approach there is substantial ambiguity in classification over which 1/3 ≤ BF ≤ 3 (where BF is computed as Probability(soft)/Probability(hard)), meaning that distinguishing between hard and soft sweeps for these paired values remains difficult or not meaningful. In addition, we find that MLGs (Figure 5) provide a greater proportion of BF ≤ 1/3 than do haplotypes (Figure S7), which yield a greater proportion of BF ≥ 3. This observation may indicate that a hard sweep with a small associated BF for MLGs will also have a small haplotype-based BF, while a hard sweep with an associated BF closer to 1, may be called as ambiguous or soft from haplotypes. We were able to address the issue of classification ambiguity with our alternative ABC approach, which assigned each test point a most probable underlying *k*. Although haplotypes provided better ability over MLGs to assign a posterior value of *k*, our results here were as expected, showing a clear increase in assigned *k* as G2/G1 or H2/H1 increased (Figure S8). For application to empirical data, however, most top sweep candidates are likely to be classifiable as hard or soft from BFs (Tables S3-S14). Pooling frequencies beyond the greatest two also increased the occupancy associated with larger BFs, and this effect was greater for haplotype data. Ultimately, the use of G123 with G2/G1 to classify sweeps and assign *k* from MLGs may be preferable because (G123, G2/G1) classification more closely resembles (H12, H2/H1) than does (G12, G2/G1). The true value of pooling additional frequencies may thus lie in sweep classification rather than detection, as G123 and H123 are not appreciably more powerful than G12 and H12 (Figures S2 and S3).

### Application of G12 and G123 to empirical data

Our analysis of human empirical data from the 1000 Genomes Project [Auton et al., 2015] recovered multiple positive controls from each study population, as well as novel candidates. Across many of these candidates, a single high-frequency MLG predominated (Figure 7). Additionally, more top candidates in CEU appear as hard sweeps than in other populations (Tables S3 and S4), though all populations had more hard sweeps than soft. The top outlying genes we detected in CEU following the application of a filter to remove heterochromatic regions with low mappability and alignability consisted of *LCT* and the adjacent loci of chromosome 2 (Figure 7A), as well as *SLC12A1* of chromosome 15 (Table S3). All of these sites are well-represented in the literature as targets of sweeps [Bersaglieri et al., 2004, Sabeti et al., 2007, Liu et al., 2013, Chen et al., 2015]. Diet-mediated selection on *LCT* likely drives the former signal cluster, as dairy farming has been a feature of European civilizations since antiquity [Itan et al., 2009, Edwards et al., 2011, Ermini et al., 2015]. Accordingly, we see that most individuals in the sample carry the most frequent MLG, and we assign this signal to be a hard sweep from its BF and from the posterior distribution of *k* generated under our demographic model for CEU (see *Materials and Methods;* Tables S3 and S4). Meanwhile, the latter signal peak is associated with the known target of selection *SLC24A5*, a melanosome solute transporter responsible for skin pigmentation [Lamason et al., 2005], also a hard sweep.

The assignment of sweeps as hard or soft in CEU, as well as their assigned *k*, were highly concordant between haplotype and MLG approaches, with the sole exception of *PRKDC*, a protein kinase involved in DNA repair [Fushan et al., 2015]. Our haplotype results indicate the presence of *k* = 3 high-frequency haplotypes at *PRKDC*, but MLG results suggest a hard sweep. This is because the window of maximum signal differs between both data types. The maximal haplotype-based window features multiple haplotypes and MLGs at high frequency, while the maximal MLG window approximately 35 kb upstream more closely resembles a hard sweep for both data types. We found such classification discrepancies to be rare across our top candidates, and typically inverted, with the MLG signal more often appearing softer (see *SYT1* and *RGS18;* Figure 7). Furthermore, we emphasize that classification discrepancies do not appear to impact the power of MLG-based methods to detect sweeps, as we generated highly concordant lists of outlying candidates for both haplotype and MLG data.

Large tracts of MLG homozygosity surround the *SYT1, RGS18, HEMGN, KIAA0825*, and *NNT* genes in YRI. Unlike for CEU, we found that assigning BFs to top signals was difficult, both for haplotype and MLG data (Tables S6 and S7). We also note a greater proportion of soft sweeps among top signals in YRI relative to other populations (Tables S6 and S7). This is likely due to the greater ease of detecting soft sweeps in more genetically diverse populations rather than any non-adaptive confounding factor (see next subsection), and we indeed see a larger occupancy of soft BFs among (G123, G2/G1) values (Figure 6). In addition, BFs for the two top candidates, *SYT1* and *RGS18*, yielded values close to 1/3 (hard) for haplotype data, but closer to 3 (soft, *k* = 2) for MLG data, indicating disproportionately large MLG diversity resulting from low haplotypic diversity, as the presence of a high-frequency haplotype alongside one or more intermediate-frequency haplotypes may generate comparatively more diversity among MLGs than haplotypes. Voight et al. [2006] previously identified our strongest selection target, *SYT1*, as a target of selection in the YRI population, and The International HapMap Consortium [2007] corroborated this, but neither speculated as to the implications of selection at this site. *SYT1* (Figure 7B) is a cell surface receptor by which the type B botulinum neurotoxin enters human neurons [Connan et al., 2017]. Selection here may be a response to pervasive foodborne bacterial contamination by *Clostridium botulinum*, similar to what exists in modern times [Chukwu et al., 2016]. Pierron et al. [2014] named *HEMGN* (which Pickrell et al. [2009] also identified), involved in erythrocyte differentiation, as a selection signal common to Malagasy populations derived from common ancestry with YRI. Racimo [2016] also identified *KIAA0825* as a target of selection, but in the ancestor to African and Eurasian populations. Our identification of *NNT* in YRI matches the result of Fagny et al. [2014], who identified this gene using a combination of iHS [Voight et al., 2006] and their derived intraallelic nucleotide diversity (DIND) method. Fagny et al. [2014] point out that *NNT* is involved in the glucocorticoid response, which is variable among global populations. Our most noteworthy candidate of selection in YRI, *RGS18*, has not been previously characterized as the location of a sweep. However, Chang et al. [2007] point to *RGS18* as a contributor to familial hypertrophic cardiomyopathy (HCM) pathogenesis. HCM is the primary cause of sudden cardiac death in American athletes [Barsheshet et al., 2011], and particularly affects African-American athletes [Maron et al., 2003].

Our scan for selection in the GIH population once again revealed the *SLC12A1* site as the strongest sweep signal (Tables S9 and S10). Because this signal is common to Indo-European populations [Liu et al., 2013, Ali et al., 2014], this was expected. However, we found that we could not confidently classify this sweep from MLG data (with inferred *k* = 2), though haplotype data suggests that this is a hard sweep. We additionally find *P4HA1* (Figure 7C) as a novel sweep candidate in GIH that exceeds the significance threshold for haplotype data, and appears as a near-soft sweep for MLGs (BF > 2.5) with inferred *k* ≥ 2 for both haplotype and MLG data. Two high-frequency MLGs predominate at the location of this candidate sweep, and their pooled frequency yields a prominent signal peak. *P4HA1* is involved in collagen biosynthesis, with functions including wound repair [Baxter et al., 2013], and the population-variable hypoxia-induced remodeling of the extracellular matrix [Petousi et al., 2013, Chakravarthi et al., 2014]. Because selection on *P4HA1* has been documented among both the tropical forest-dwelling African pygmy population [Mendizabal et al., 2012, Amorim et al., 2015] and now in individuals of Gujarati descent, and is known to present a differing expression profile among low- and high-altitude populations [Petousi et al., 2013], this gene may be involved in a number of adaptations to harsh climatic conditions, potentially in wound repair, which is more difficult in tropical climates.

Of the sweep candidates we identified in the CHB population (Tables S12 and S13), we found that the inferrence of significance from G123 was considerably more concordant with H12 than was G12. We recovered as top candidates *EXOC6B*, which produces a protein component of the exocyst [Evers et al., 2014] and *LONP2*, both previously documented [Baye et al., 2009, Ohashi et al., 2010, Durbin and Consortium, 2011, Pybus et al., 2014]. *EXOC6B* is a characteristic signal in East Asian populations alongside *EDAR*, which we did not specifically recover in our scan (but nearby candidates *LIMS1, CCDC138*, and *RANBP2* did appear), while *LONP2* is adjacent to *ABCC11*, which controls earwax texture. *FMNL3* yielded elevated values of G12 and G123 in CHB, but was only significant from its H12 value. A single MLG predominates at *FMNL3* in the sample (Figure 7D), and all approaches assign this sweep as hard. The function of *FMNL3* is related to actin polymerization [Hetheridge et al., 2012, Gauvin et al., 2014], and has a role in shaping the cytoskeleton, which it shares with *EXOC6B*. Moreover, the signal at *FMNL3* may be additionally associated with the outlier *RANBP10*, which also interacts with the cytoskeleton, but with microtubules [Schulze et al., 2008]. Though it is unclear why we identify an enrichment in cytoskeleton-associated genes, future studies may shed light on why variants in such genes could be phenotypically-relevant specifically in individuals of East Asian descent. Finally, we found *SPATA31D3* as a hard sweep within the top H12 signals in CHB, as well as in GIH, and while it did not exceed our significance threshold, this is in line with the results of Schrider and Kern [2017].

### Addressing confounding scenarios

A variety of processes, both adaptive and non-adaptive, may produce elevated values of expected homozygosity in the absence of selective sweeps in a sampled population, or small values of expected homozygosity despite a sweep, thereby misleading expected homozygosity methods. To understand the impacts of potentially confounding processes on the power of the expected homozygosity methods, we evaluated the effects of long-term background selection, long-term population substructure, and pulse admixture on G12, G123, H12, and H123. We additionally consider the confounding effect of missing data, as the manner in which missing sites is addressed during computations can change analyzed patterns of MLG and haplotype diversity.

We first addressed long-term background selection as a potentially common confounding factor with a brief experiment to determine the susceptibility of all methods to the misidentification of background selection as a sweep. Signatures of background selection are ubiquitous in a number of systems [McVicker et al., 2009, Comeron, 2014], and the effect of background selection is a reduction in nucleotide diversity and a distortion of the site frequency spectrum, which to many methods may spuriously resemble a sweep [Charlesworth et al., 1993, 1995, Seger et al., 2010, Charlesworth, 2012, Nicolaisen and Desai, 2013, Cutter and Payseur, 2013, Huber et al., 2016]. Here, we simulated chromosomes containing a centrally-located genic region of length 11 kb in which deleterious alleles arise throughout the course of the simulation. Our model involved a gene with exons, introns, and untranslated regions (UTRs) with properties based on human-inspired parameters (see *Materials and methods)*. In agreement with the result of Enard et al. [2014], we found that background selection did not distort the haplotype (and therefore MLG) frequency spectrum to resemble that of a sweep, such that G12 and G123 were thoroughly robust to background selection. We demonstrate this by displaying the concordance in the distributions of maximum G12, G123, H12, and H123 scores for background selection and neutral evolution scenarios (Figure S20). Thus, we do not expect that outlying G12, G123, H12, or H123 values can result from background selection.

Methods to detect recent sweeps may be confounded by the effect of long-term population substructure, as well as from admixture. Structured populations contain a greater proportion of homozygous genotypes than would be expected under an equally-sized, randomly-mating population [Sinnock, 1975], thereby increasing the chance that an elevated level of expected homozygosity will arise in the absence of a sweep. We examined the possibility that a symmetric island migration model with six demes (Figure S21A), and migration rates (*m*) between demes of *m* ∈ {10^−5^, 10^−4^, 10^−3^, 10^−2^, 10^−1^} per generation (a proportion *m* of the haplotypes in a deme derives from each of the other five demes for a total proportion of 5*m* haplotypes) could yield elevated values of H12 and G123 under neutrality. We found that compared to a model with no substructure, H12 and G123 values were moderately impacted for a model with population substructure. These values were substanially lower than expected H12 and G123 values under a recent strong hard sweep. However, these values are more comparable to an ancient sweep, and so caution is warranted in the study of structured populations for all but the most outlying signals.

The expected homozygosity methods are similarly robust to the effect of admixture under most scenarios. Specifically, we evaluated whether any admixture scenario can falsely generate a signature of a sweep. We simulated a model in which a single ancestral population diverges into two descendant populations (Figure S21B; see also *Materials and methods*). We maintained the size of one descendant population (the target) at *N* = 10^4^ diploid individuals, and varied the size of the unsampled (donor) population (*N* = 10^3^, 10^4^, or 10^5^ diploids), admixing at rate *m G* {0.05, 0.10,0.15,0.20, 0.25, 0.30,0.35,0.40} as a single pulse 200 generations before sampling. We find that admixture with donor sizes *N* = 10^5^ or *N* = 10^4^ produces only small values of H12 (Figure S23, left and center) and G123 (Figure S24, left and center) in the sampled population in the absence of a sweep. However, admixture with a donor population of small size (*N* = 10^3^) can produce elevated values of H12 and H2/H1, as well as G123 and G2/G1 when migration is sufficiently large (*m* ≥ 0.15), thus spuriously resembling the pattern of a soft sweep in the absence of selection (Figures S23 and S24, right). In this scenario, with a large enough admixture fraction, there will be a high probability that many sampled lineages from the target population will derive from the donor population, which will coalesce rapidly due to the small effective size, which will in turn lead to elevated homozygosity. Small donor population sizes with large migration rates therefore represent the only admixture scenario that we considered under which the expected homozygosity methods are susceptible to misclassifying neutrality as selection, specifically as a soft sweep. Otherwise, our methodology remains robust under a wide range of other admixture scenarios. We note therefore that the elevated number of soft sweeps we detected within the YRI population (Tables S6 and S7) is unlikely to be due to the effect of the admixture described in Busby et al. [2016], as this would produce a genome-wide pattern, which we do not observe (Figures S13 and S14).

Finally, we note that accounting for missing data is a practical consideration that must be undertaken when searching for signals of selection, and the manner in which missing data are removed affects our ability to identify sweeps. We explored the effects of two corrective strategies to account for missing data. Our strategies were to remove sites with missing data or to define MLGs and haplotypes with missing data as new distinct MLGs and haplotypes. Relative to the ideal of no missing data (Figure 3A), removing sites resulted in a slight inflation of power observed in the absence of missing data. This was true for G12 and H12 (Figure S25A), as well as G123 and H123 (Figure S25C). After removing sites, the overall polymorphism in the sample decreases, but windows containing the site of selection are still likely to be the least polymorphic, and therefore identifiable. Even so, weaker sweeps are likely to be obscured by the lower background diversity after removing sites. Conservatively defining MLGs and haplotypes with missing data as new distinct MLGs and haplotypes inflates the total observed diversity and results in a more rapid decay of power compared to complete data (Figures S25B and D). This result is because individuals affected by the sweep may have different patterns in their missing data, and therefore different assigned sequences after accounting for missingness. Overall, the choice of strategy will likely depend on the level of missing data in the sample. Removing too many sites is likely to generate false positive signals, while removing no sites may lead to false negatives.

### Concluding remarks

Our results emphasize that detecting selective sweeps does not require phased haplotype data, as distortions in the frequency spectrum of MLGs capture the reduction in diversity under a sweep similarly well to phased haplotypes. Accordingly, the advent of rapid and cost-effective genotyping-by-sequencing technologies [Elshire et al., 2011] across diverse taxa including bovine, marine-dwelling, and avian populations means that the adaptive histories of myriad organisms may now be inferred from genome-wide data [Daetwyler et al., 2014, Drury et al., 2011, Zhu et al., 2016]. Furthermore, we have shown that the inferences emerging from MLG-based scans align with those of phased haplotype-based scans, with empirical analyses of human populations yielding concordant top outlying candidates for selection, both documented and novel. We demonstrate as well that paired (G12, G2/G1) and (G123, G2/G1) values properly distinguish hard sweeps from soft sweeps. In addition to identifying sweeps from single large values of G12 and G123, we find that the genomic signature of these MLG-based statistics surrounding the site of selection provides a means of distinguishing a sweep from other types of selection (e.g., balancing selection). This additional layer of differentiation motivates the use of MLG identity statistics as a signature in a statistical learning framework, as such approaches have increasing in prominence for genome analysis [Grossman et al., 2010, Lin et al., 2011, Pavlidis et al., 2010, Ronen et al., 2013, Pybus et al., 2015, Ronen et al., 2015, Sheehan and Song, 2016, Schrider and Kern, 2016, Akbari et al., 2017, Kern and Schrider, 2018, Mughal and DeGiorgio, 2018]. We expect that the MLG-based approaches G12 and G123, in conjunction with G2/G1, will be invaluable in localizing and classifying adaptive targets in both model and non-model study systems.

## Materials and methods

### Simulation parameters

To compare the powers of G12 and G123 to detect sweeps relative to H12 and H123 [Garud et al., 2015], we performed simulations for neutral and selection scenarios using SLiM 2 (version 2.6) [Haller and Messer, 2017]. SLiM is a general-purpose forward-time simulator that models a population according to Wright-Fisher dynamics [Fisher, 1930, Wright, 1931, Hartl and Clark, 2007] and can simulate complex population structure, selection events, recombination, and demographic histories. For our present work, we used SLiM 2 to model scenarios of recent selective sweeps, long-term background selection, and neutrality, additionally including models of population substructure and pulse admixture. Our models of sweeps comprised complete and partial hard sweeps, as well as soft sweeps from selection on standing variation (SSV). For background selection, we simulated a gene with introns, exons, and untranslated regions in which deleterious mutations arose randomly. We additionally tested the effect of demographic history on power by examining constant population size, population expansion, and population bottleneck models for hard sweep scenarios.

### General approach

We first simulated data according to human-specific parameters for a constant population size model. For simulated sequences (Figures 2A and D), we chose a mutation rate of *μ* = 2.5 × 10^−8^ per site per generation, a recombination rate of *r* = 10^−8^ per site per generation, and a diploid population size of *N* = 10^4^ [Takahata et al., 1995, Nachman and Crowell, 2000, Payseur and Nachman, 2000]. All simulations ran for a duration of 12*N* generations, where *N* is the starting population size for a simulation, equal to the diploid effective population size. The duration of simulations is the sum of a 10N generation burn-in period of neutral evolution to generate equilibrium levels of variation across simulated individuals [Messer, 2013], and the expected time to coalescence for two lineages of 2*N* generations. Simulation parameters were scaled, as is common practice, to reduce runtime while maintaining expected levels of population-genetic variation, such that mutation and recombination rates were multiplied by a factor λ, while population size and simulation duration were divided by λ. For simulations of constant population size, we used λ = 20.

Scenarios involving population expansion and bottleneck were modeled on the demographic histories inferred by Lohmueller et al. [2009]. For population expansion (Figures 2B and D), we used λ = 20, and implemented the expansion at 1,920 unscaled generations before the simulation end time. After expansion, the size of the simulated population doubled from 10^4^ to 2 × 10^4^ diploid individuals. This growth in size corresponds to the increase in effective size of African populations that occurred approximately 48,000 years ago [Lohmueller et al., 2009], assuming a generation time of 25 years. Population bottleneck simulations (Figures 2C and D) were scaled by λ = 10, began at 1,200 generations before the simulation end time, and ended at 880 generations before the simulation end time. During the bottleneck, population size fell to 550 diploid individuals. This drop represents the approximately 8,000-year bottleneck that the population ancestral to non-African humans experienced as it migrated out of Africa [Lohmueller et al., 2009], assuming a generation time of 25 years.

### Simulating selection

Our simulated selection scenarios encompassed a variety of selection modes and parameters. Though we primarily focused on selective sweeps, we additionally modeled a history of long-term background selection to test the specificity of methods for sweeps. Background selection may decrease genetic diversity relative to neutrality. For sweep experiments specifically, we tested the power of methods to detect selection occurring between 40 and 4,000 generations prior to the simulation end time (thus, within 2*N* generations prior to sampling). We set the site of selection to be at the center of the simulated chromosome, and performed two categories of simulations, allowing us to answer two distinct types of questions about the power of our approach: whether G12 and G123 properly identify the signature of a selective sweep (the detection experiments), and whether G12 or G123 in conjunction with G2/G1 can distinguish between hard and soft sweeps and ultimately infer the number of selected haplotypes (*k*; the classification experiments), and hence “‘softness”’ of the sweep.

For the detection experiments (see *Detecting sweeps*), we simulated chromosomes of length 100 kb under neutrality and for each set of selection parameters, performed 10^3^ replicates of sample size *n* = 100 diploids (and *n* = 25 for hard sweep experiments in Figure S19). Here, we fixed the times (*t*) at which selected alleles arise to be 400, 1,000, 2,000, or 4,000 generations prior to sampling (Figure 2), and selection coefficients (*s*) to be either 0.1 or 0.01, respectively representing strong and moderate selection. The parameters *t* and *s* were common to all selection simulations of the first type, with additional scenario-specific parameters which we subsequently define. For the classification experiments (see *Differentiating between hard and soft sweeps*), we performed two types of simulations. First, we simulated 10^6^ replicates of *n* = 100 diploids for each scenario, with *s* ∈ [0.005,0.5], drawn uniformly at random from a natural log-scale, and *t* ∈ [40, 2000] (also drawn uniformly at random from a natural log-scale), across chromosomes of length 40 kb. With these simulations, we assessed the occupancy of specific hard and soft sweeps among (G12, G2/G1), (G123, G2/G1), (H12, H2/H1), and (H123, H2/H1) test points. Second, we simulated 5 × 10^6^ replicates with *s* ∈ [0.05,0.5] and *t* ∈ [200, 2000] and all other parameters as previously. Here, we assigned the most probable *k* to each test point from the posterior distribution of *k* among nearby test points, drawing *k* ∈ {1, 2, …, 16} uniformly at random. We scaled selection simulations as previously described.

We first examined hard sweeps, in which the beneficial mutation was added to one randomly-drawn haplotype from the population at time *t*, remaining selectively advantageous until reaching a simulation-specified sweep frequency (*f*) between 0.1 and 1.0 at intervals of 0.1, where *f* < 1.0 represents a partial sweep and *f* = 1.0 is a complete sweep (to fixation of the selected allele). Although we conditioned on the selected allele not being lost during the simulation, we did not require the selected allele to reach *f*. We additionally modeled soft sweeps from selection on standing genetic variation (SSV). For this scenario, we introduced the selected mutation to multiple different, but not necessarily distinct, randomly-drawn haplotypes (*k*) such that *k* = 2, 4, 8, 16, or 32 haplotypes out of 2*N* = 10^3^ (scaled haploid population size) acquired the mutation at the time of selection. We did not condition on the number of remaining selected haplotypes at the time of sampling as long as the selected mutation was not lost.

For hard sweeps only, we additionally examined the effects of three common scenarios—population substructure, pulse admixture, and missing data—on performance. The population substructure model consisted of six demes in a symmetric island migration model in which migration between each deme is constant at rate *m* per generation for the duration of the simulation (Figure S21A). We simulated *m* ∈ {10^−5^, 10^−4^, 10^−3^, 10^−2^, 10^−1^}. All demes were identical in size at *N* = 1, 660 (unscaled) diploid individuals, and samples consisted of *n* = 100 diploid individuals, with 50 individuals sampled from each of two demes. Thus, as *m* increases, the structured model converges to the unstructured model of *N* = 10^4^ (unscaled) diploid individuals. Our admixture scenarios examined a single pulse of gene flow from an unsampled donor population into the sampled target at rate *m* ∈ {0.05,0.10, 0.15, 0.20,0.25,0.30, 0.35, 0.40}, occurring 200 generations prior to sampling (Figure S21B). We performed experiments in which the donor had a (unscaled) diploid size of *N* = 10^3^, 10^4^, or 10^5^, keeping the size of the target fixed at *N* = 10^4^. For admixture simulations, a single population of size 10^4^ diploids evolves neutrally until it splits into two subpopulations at 4, 000 generations before sampling. We selected the divergence and admixture times to approximately match the timing of these events in sub-Saharan African populations [Veeramah et al., 2011, Busby et al., 2016]. Sample sizes were of *n* = 100 diploids, matching the standard hard sweep experiments.

To simulate missing data in the sampled population, we followed a random approach. Using data generated for the previous simple hard sweep experiment, we removed data from a random number of SNPs in each replicate sample, between 25 and 50, drawing these sites from locations throughout the simulated sequence uniformly at random. At each missing site, we assigned a number of the sampled individuals, between 1 and 5, uniformly at random, to have their genotypes missing at the site. We then accounted for missing data in one of two ways. First, we omitted any SNP with missing data in each analysis window. This reduced the number of SNPs included in each computation. Second, we assigned any haplotype or MLG with missing data as an entirely new string. Thus, the number of distinct haplotypes and MLGs increases when sites are missing, providing a more conservative approach than the first.

Finally, our single scenario of background selection was intended to quantify the extent to which the long-term removal of deleterious alleles in a population, which reduces nearby neutral genetic diversity, would mislead each method to make false inferences of selective sweeps. We generated a 100 kb chromosome containing an 11 kb gene at its center and allowed it to evolve over 12N generations under a constant-size demographic model. The gene was composed of 10 exons of length 100 bases with 1 kb introns separating each adjacent exon pair. The first and last exons were flanked by untranslated regions (UTRs) of length 200 bases at the 5’ end and 800 bases at the 3’ end. Strongly deleterious mutations (*s* = − 0.1) arose at a rate of 50% in the UTRs, 75% in exons, and 10% in introns, while mutations occurring outside of the genic region were neutral. To measure the confounding effect of background selection, we observed the overlap between the distributions of maximum G12, G123, H12, and H123 values of 10^3^ simulated replicates under neutrality and background selection. Our model here is identical to that of Cheng et al. [2017], with the sizes of genetic elements based on human mean values [Mignone et al., 2002, Sakharkar et al., 2004].

### Detecting sweeps

We performed scans across simulated 100 kb and one Mb chromosomes with all methods using sliding genomic windows of length 40 kb, advancing by four kb increments. We chose this window size primarily because the mean value of LD between pairs of loci across the chromosome decays below one-third of its maximum value over this interval (Figure S1), and because this was the window size with which we analyzed all non-African populations from the 1000 Genomes dataset. Window size also affects sensitivity to sweeps by constraining the minimum strength of selective sweeps we can detect. That is, with our chosen window size, we are likely to detect sweeps with *s* > 0.004, because such sweeps will generate genomic footprints on the order of 40 kb for our simulated population size of *N* = 10^4^. We computed this value as *F* = *s*/(2*r* ln(4Ns)), where *F* is the size of the footprint in nucleotides, *s* is the per-generation selection coefficient, *r* is the per-base, per-generation recombination rate, and *N* is the effective population size [Gillespie, 2004, Garud et al., 2015, Hermisson and Pennings, 2017].

For experiments measuring power at defined time points, we recorded the chromosomal maximum value of G12, G123, H12, or H123 across all windows as the score for each of 10^3^ replicates of 100 kb chromosomes. Selection simulation scores provided us with a distribution of values that we compared with the distribution of scores generated under neutral parameters. We define a method’s power for each of our specified time intervals at the 1% false positive rate (FPR). This measures the proportion of our 1,000 replicates generated under selection parameters with a score greater than the top 1% of scores from the neutral replicates. The method performs ideally if the distribution of its scores under a sweep does not overlap the distribution of scores for neutral simulations; *i.e.*, if neutrality can never produce scores as large as a sweep.

In addition to power, we also tracked the mean scores of G12 and G123 across simulated one Mb chromosomes at each 40 kb window for all selection scenarios at the time point for which power was greatest. In situations where G12 or G123 had the same power at more than one time point (this occurred for strong selection within 1,000 generations of sampling), we selected the most recent time point in order to represent the maximum signal, since mutation and recombination erode expected haplotype homozygosity over time. This analysis allowed us to observe the interval over which elevated scores are expected, and additionally define the shape of the sweep signal.

### Differentiating between selection scenarios

Experiments to test the ability of G2/G1 to correctly differentiate between soft and hard sweeps, as H2/H1 can (conditioning on a G12 or G123 value for G2/G1, or an H12 or H123 value for H2/H1), required a different simulation approach than did the simple detection of selective sweeps. Whereas multiple methods exist to identify sweeps from extended tracts of expected haplotype homozygosity, the method of Garud et al. [2015] classifies this signal further to identify it as deriving from a soft or hard sweep. As did Garud et al. [2015], we undertook an approximate Bayesian computation (ABC) approach to test the ability of our method to distinguish soft and hard sweeps. To demonstrate the ability of G2/G1 conditional on G12 and G123 to differentiate between sweep scenarios and establish the basic properties of the (G12, G2/G1) and (G123, G2/G1) distributions, we simulated sequences of length 40 kb under a constant population size demographic history (Figure 2A) with a centrally-located site of selection. Here, we treated the whole simulated sequence as a single window.

For ABC experiments to classify test points as hard or soft from a fixed number of different selected haplotypes *k*, we performed 10^6^ simulations for each selection scenario, drawing selection coefficients *s* and selection times *t* uniformly at random from a log-scale as previously described. Soft sweeps from SSV were generated for *k* = 5 and *k* = 3 starting haplotypes (out of a scaled 2*N* = 10^3^ haploids). Soft sweeps generated under random *t* and *s* were compared with hard sweeps generated under random *t* and *s*, with completion of the sweep possible but not guaranteed. From the resulting distribution of scores for each simulation type, we computed Bayes factors (BFs) for direct comparisons between a hard sweep scenario and either soft sweep scenario.

For two selection scenarios *A* and *B* and a (G12, G2/G1) or (G123, G2/G1) test point (or haplotype statistic test point), we compute BFs as the number of simulations of type *A* yielding results within a Euclidean distance of 0.1 from the test point, divided by the number of simulations of type *B* within that distance. Here, test values of (G12, G2/G1) and (G123, G2/G1) are each plotted as a 100 × 100 grid, with both dimensions bounded by [0.005,0.995] at increments of 0.01. In the work of Garud et al. [2015], soft sweeps were of type *A* and hard sweeps were of type B, and we retain this orientation in our present work. Following these definitions, a BF less than one at a test coordinate indicates that a hard sweep is more likely to generate such a (G12, G2/G1) or (G123, G2/G1) pair, whereas a BF larger than one indicates greater support for a recent soft sweep generating that value pair. As do Lee and Wagenmakers [2013], we define BF ≥ 3 as representing evidence for selection scenario *A* producing a similar paired (G12, G2/G1) or (G123, G2/G1) value as the test point, and BF ≥ 10 to represent strong evidence. Similarly, BF ≤ 1/3 is evidence in favor of scenario B, and BF ≤ 1/10 is strong evidence. We performed analyses for both MLG and haplotype data to demonstrate the effect of data type on sweep type inference.

We followed a similar approach for ABC experiments to assign a most probable *k* to test points within the aforementioned 100 × 100 grids. Here, we generated 5 × 10^6^ replicates, drawing *t* and *s* uniformly at random on a log scale as previously, and *k* ∈ {1, 2, …, 16} uniformly at random. For each (G12, G2/G1), (G123, G2/G1), (H12, H2/H1), or (H123, H2/H1) test point, we retained the value of *k* for each replicate within a Euclidean distance of 0.1, and assigned the most frequently-occurring *k* as the most probable value for the test point. Thus, unlike for BF experiments, no test point yielded an ambiguous result, and all test points were assigned a most probable *k*.

### Analysis of empirical data

We evaluated the ability of G12, G123, and H12 to corroborate and complement the results of existing analyses on human data. Because G12 and G123 take unphased diploid MLGs as input, we manually merged pairs of haplotype strings for this dataset (1000 Genomes Project, Phase 3 [Auton et al., 2015]) into MLGs, merging haplotype pairs that belonged to the same individual. We also complemented the individual-centered approach by randomly merging pairs of haplotypes to produce a sample of individuals that could arise under random mating. Our approaches therefore allowed us to determine the effect of using different data types to infer selection. Unlike biallelic haplotypes, MLGs are triallelic, with an indicator for each homozygous state and the heterozygous state. Thus, there are at least as many possible MLGs as haplotypes, such that a sample with *I* distinct haplotypes can produce up to *I* (*I* + 1)/2 distinct MLGs.

We scanned all autosomes using nucleotide-delimited genomic windows, proportional to the effective size of the study population, and the interval over which the rate of decay in pairwise LD plateaus empirically [see Jakobsson et al., 2008]. For the 1000 Genomes YRI population, we employed a window of length 20 kb sliding by increments of two kb, whereas for non-African populations (effective population size approximately half of YRI) we used a window of 40 kb sliding by increments of five kb (see *Results*). This means that we were sensitive to sweeps from approximately *s* ≥ 0.002 for YRI, and approximately *s* ≥ 0.004 for the others. We recorded G12, G123, and H12 scores for all genomic windows, and subsequently filtered windows for which the observed number of SNPs was less than a certain threshold value in order to avoid biasing our results with heterochromatic regions for which sequence diversity is low in the absence of a sweep. Specifically, we removed windows containing fewer SNPs than would be expected [Watterson, 1975] when two lineages are sampled, which is the extreme case in which the selected allele has swept across all haplotypes except for one. For our chosen genomic windows and all populations, this value is 4*N_e_μ* × (window size in nucleotides) = 40 SNPs, where *N_e_* is the diploid effective population size and *μ* is the per-site per-generation mutation rate. As in Huber et al. [2016], we additionally divided each chromosome into non-overlapping 100 kb bins and removed sites within bins whose mean CRG100 score [Derrien et al., 2012], a measure of site mappability and alignability, was less than 0.9. Filtering thereby removed additional sites for which variant calls were unreliable, making no distinction between genic and non-genic regions.

Following a scan, we intersected selection signal peaks with the coordinates for protein- and RNA-coding genes and generated a ranked list of all genomic hits discovered in the scan for each population. We used the coordinates for human genome build hg19 for our data, to which Phase 3 of the 1000 Genomes Project is mapped. The top 40 candidates for each study population were recorded and assigned *p*-values and BFs. Specifically, we simulated sequences following the estimates of population size generated by Terhorst et al. [2017] from `smc++` using *ms* [Hudson, 2002] to assign *p*-values and SLiM 2 to assign BFs, with per-generation, per-site mutation and recombination rates of 1.25 × 10^−8^ and 3.125 × 10^−9^ [Terhorst et al., 2017, Narasimhan et al., 2017], and sample sizes for each population matching those of the 1000 Genomes Project. For *p*-value simulations, we selected a sequence length uniformly at random from the set of all hg19 gene lengths, appended the window size used for scanning that population’s empirical data to this sequence, and used a sliding window approach, retaining information from the window of maximum G12, G123, or H12 value. For BF simulations, we used simulated sequence lengths of either 20 kb for YRI or 40 kb for others, to match the strategy of empirical scans. That is, once we have identified an elevated sweep signal within a window, we then seek to classify it as hard or soft.

We assigned *p*-values by generating 10^6^ replicates of neutrally-evolving sequences, where the *p*-value for a gene is the proportion of maximum G12 (or G123 or H12) scores generated under neutrality that is greater than the score assigned to that gene. After Bonferroni correction for multiple testing [Neyman and Pearson, 1928], a significant *p*-value was *p* < 0.05/23, 735 ≈ 2.10659 × 10^−6^, where 23,735 is the number of protein- and RNA-coding genes for which we assigned a G12 (or G123 or H12) score. To assign BFs, we simulated 10^6^ replicates of hard sweep and SSV (*k* = 5) scenarios for each study population (thus, 2 × 10^6^ replicates for each population), wherein the site of selection was at the center of the sequence. We drew *t* ∈ [40, 2000] and *s* ∈ [0.005,0.5] uniformly at random from a log-scale, and defined BFs as previously. Additionally, we assigned the most probable values of *k* from the posterior distribution for each top 40 sweep candidate for each population, following the previous protocol. Values of *t* were chosen to reflect selective events within the range of detection of G12, G123, and H12, while also being after the out-of-Africa event, whereas values of *s* represent a range of selection strengths from weak to strong. We once again conditioned on the selected allele remaining in the population throughout the simulation, though not on its frequency beyond this constraint.

We affirm that all data necessary for confirming the conclusions of the article are present within the article, figures, and tables. Any other materials and resources are available upon request.

## Acknowledgments

This work was supported by National Institutes of Health grant R35GM128590, by the Alfred P. Sloan Foundation, and by Pennsylvania State University startup funds. We also thank Jonathan Terhorst for providing demographic information on our study populations, estimated from his method `smc++`, as well as Dmitri Petrov, Pleuni Pennings, and Arbel Harpak for helpful conversations. Finally, we thank three anonymous reviewers for evaluating the merit of this work and providing comments that improved its overall quality. Portions of this research were conducted with Advanced CyberInfrastructure computational resources provided by the Institute for CyberScience at Pennsylvania State University.

